# Global analysis of trimeric autotransporters reveals phylogenetically restricted secretion mechanism adaptations

**DOI:** 10.64898/2026.01.22.700529

**Authors:** Louis Dorison, Bianca Audrain, Susan Chamorro-Rodriguez, Jean-Marc Ghigo, Christophe Beloin

## Abstract

Autotransporters are important diderm bacterial cell-surface proteins and are virulence factors enabling surface attachment and adhesion to other bacteria. These proteins are composed of a signal peptide, a β-barrel that serves as an anchor in the outer membrane, and an extracellular passenger domain responsible for adhesion. Autotransporters rely on BamA for their insertion in the outer membrane (OM), but specific helper proteins, such as TpgA and SadB have been described to promote the surface exposure of trimeric autotransporters (TAAs), a specialized subclass of autotransporters forming homotrimer adhesins. To identify domains or proteins that could help TAAs secretion, we analyzed a recent dataset of all trimeric autotransporters found across the bacterial tree of life. While we did not find additional potential helper proteins for OM translocation, we found that the extended signal peptide (ESPR), sometimes found in TAAs, is associated with longer adhesins both for TAAs and type Va autotransporter adhesins. ESPRs are found in all bacteria but Fusobacteriia and Alphaproteobacteria. We also identified in *Burkholderia*, *Veillonellales* and *Pasteurellales* a DUF2827 domain proteins as potential glycosyltransferases constantly associated with TAAs. Finally, e describe the existence of extra periplasmic domains in some TAAs, featuring either a coiled-coil domain or a peptidoglycan-binding domain. Our research show that there is a strong phylogenetic separation between Terrabacteria, almost invariably displaying additional periplasmic domains, and Gracilicutes (represented by Proteobacteria) where they are largely absent. This suggests that the presence these domains might be correlated with specific Terrabacteria OM features. Using the diderm Firmicute *Veillonella parvula* as a model, we demonstrate that the absence of periplasmic domains in TAAs leads to a significant protein degradation, yet they are not essential for adhesin trimerization or secretion. Additionnaly, we show that the SLH domains of *V. parvula* TAAs excludes them from the septum during division, but that this exclusion is not crucial for adhesin function or stability in the tested conditions. Altogether, these results illuminate the genetic flexibility and modularity of autotransporters, enhancing our understanding of this important class of adhesins in diderm bacteria.

## Introduction

Diderm (Gram-negative) bacteria often need to translocate proteins and effectors through their outer membrane, but their periplasm lacks ATP for active transport. To address this, diderms have evolved secretion systems (SS) such as the type I-IV and type VI secretion systems, which form large transmembrane complexes, and the type V secretion system, which relies on the classical outer membrane protein assembly pathway. Proteins of the type V secretion system, also known as autotransporters, are composed of a single protein encoding a signal peptide for inner membrane translocation via the SEC machinery and a hydrophobic β-barrel inserted into the outer membrane by the BAM machinery that functions as an anchor for the passenger (or effector) domain, sometimes connected by a stalk domain (1). In rare cases, the TAM machinery has been linked to autotransporter secretion (2–4) Autotransporters vary in the size of their β-barrel, the type of passenger domain, and the presence of a stalk.

Autotransporters often have additional characteristics or factors that aid their secretion. Some possess an extended signal peptide region (ESPR) of 50-60 residues, thought to slow secretion through the SEC pathway and prevent misfolding in the periplasm (5). Trimeric autotransporters (TAAs), which correspond to the Type Vc SS, form homotrimeric adhesins where each protein contributes four β-strands to a twelve stranded β-barrel (6). Some helper proteins for TAA secretion have been identified; in *Salmonella*, the inner membrane trimeric lipoprotein SadB promotes the surface exposure of the SadA TAA (7). TpgA, a periplasmic protein interacting with the TAA AtaA from *Acinetobacter* facilitates the passenger domain secretion but is not crucial for insertion in the outer membrane (8). TpgA comprises a BamE-like domain that likely binds directly to the AtaA β-barrel and an OmpA peptidoglycan-binding domain. TpgA is potentially important for anchoring the TAA to the cell (9). Moreover, inverted autotransporters (type Ve SS) often have periplasmic domains, either a short coiled-coil domain or a LysM peptidoglycan-binding domain. Deleting the LysM domain in YeeJ *of Escherichia coli* reduces surface exposure of the adhesin (10). This LysM domain can bind peptidoglycan and mediate autotransporter dimerization, whereas the coiled-coil domain cannot perform these functions (11).

We recently observed that similarly to inverted autotransporters periplasmic domains, a subset of trimeric autotransporters possesses a periplasmic extension at their C-terminus, right after their β-barrel, characterized by either a coiled-coil domain or a S-layer homology SLH peptidoglycan-binding domain (12). We wondered whether these extensions were more widespread in bacteria and if TAAs could possess additional helper proteins or domains previously undetected. We first used bioinformatic analysis on a dataset of all TAAs sequences identified in UniProt to identify additional secretion factors. Using the Negativicutes *Veillonella parvula*, a diderm firmicute rich in TAAs, we then investigated the functions of these previously undescribed periplasmic domains.

## Results

### Extended Signal peptide Regions (ESPR) are associated with longer autotransporters and are phylogenetically restricted

To identify potential additional domains important for secretion present in TAAs, we generated an exhaustive database of TAAs sequences by parsing the Uniprot database for all YadA-anchor domain containing proteins (IPR005594), which resulted in 12394 unique protein sequences, with 7193 of them also including a YadA head (IPR008640) and YadA stalk domains (IPR008635). These are predominantly found in Gamma- (6655 TAAs), Beta- (2934 TAAs) and Alphaproteobacteria (1104 TAAs), followed by Negativicutes (1021 TAAs) and Fusobacteriia (345 TAAs), with a notable overrepresentation of frequent pathogens such as *Escherichia* and *Salmonella* (supplementary file S1). Using this database, we started by investigating the presence of ESPR, a domain known to slow down autotransporter secretion through the Sec translocon (13,14). Across all phylogenetic classes, TAAs containing an ESPR are, on average, twice as long as those lacking it. Specifically, TAAs with ESPR exhibit passenger domains of approximately 2000 amino acid residues, compared to about 1000 residues in those without ESPR (Figure 1). Interestingly, we observed a similar trend in classical autotransporters (Va class) (supplementary Figure S1), where the presence of ESPR also correlates with longer passenger domains - around 2000 residues versus 1000 in its absence. We noticed that *Alphaproteobacteria* (already identified in (15)) and *Fusobacteriia* TAAs are characterized by a lack of ESPR (Figure 1), which is also the case for their monomeric autotransporters (supplementary Figure S1). The predicted size of their signal peptides typically matched the usual length, with no increased prevalence of longer signal peptides that might suggest an alternative, undetected type of ESPR signal peptide (supplementary Figure S2).

**Figure 1:**
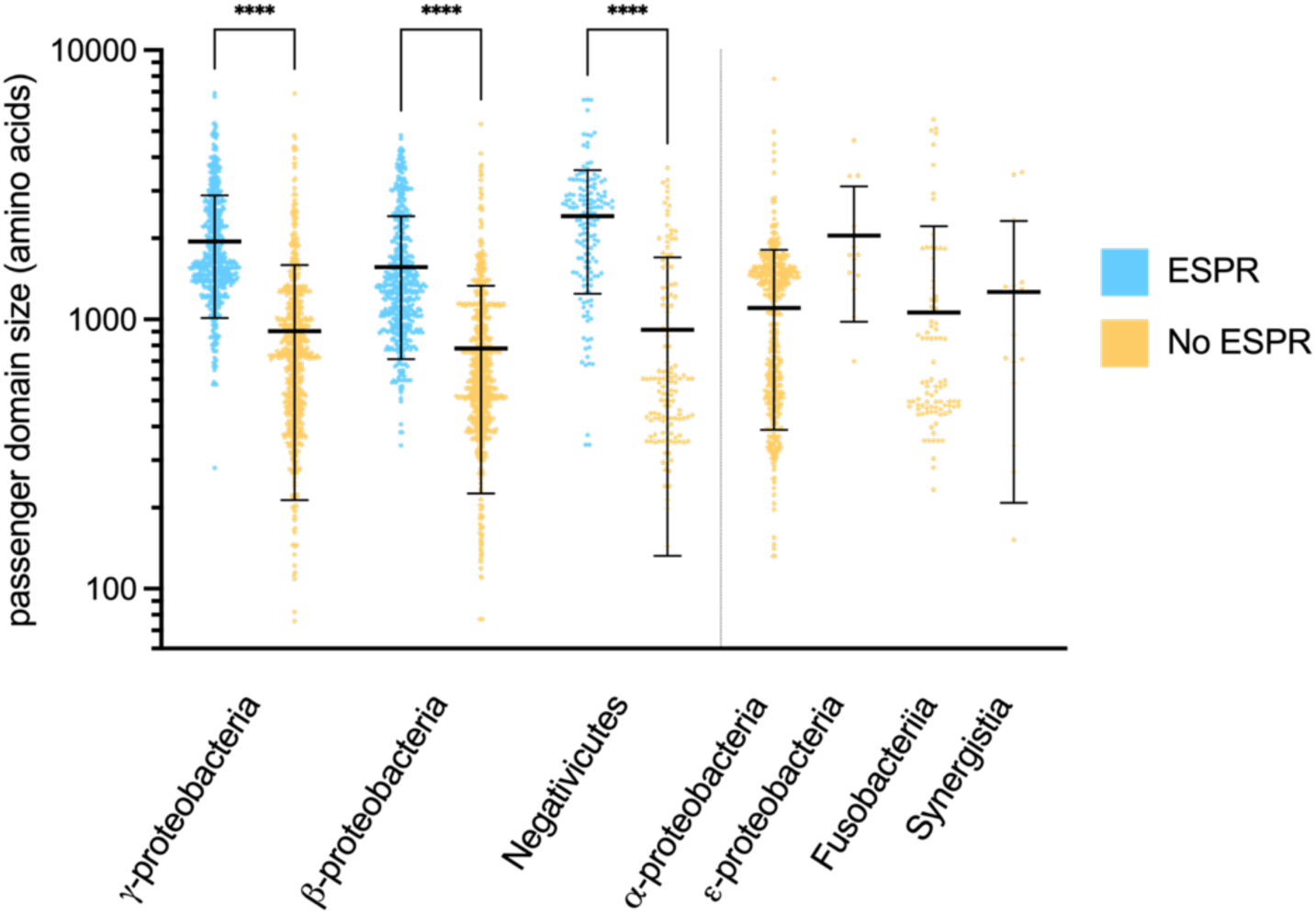
TAAs with ESPR are on average longer than adhesins without ESPR. Scatter plot of TAAs, where each point corresponds to the size in amino acid of their passenger domain. Mean and standard deviation are also represented. TAAs were separated by Class and presence or absence of ESPR. Significance of the differences between adhesins with and without ESPR was calculated using a Kruskal-Wallis test (**** : P ≤ 0.0001).

### Search for additional TAA helper proteins

TpgA and SadB are the only described TAAs-associated proteins that faciliate TAAs secretion, and we wondered if more TAAs could rely on a larger diversity of helper proteins. For this, we retrieved from our TAA database 1383 genomes (supplementary file S2), each containing at least one trimeric autotransporter with both a YadA anchor (IPR005594) and a YadA head domain (IPR008640). With a “guilty by association” hypothesis, we extracted and clustered the protein sequences of all TAAs adjacent genes (+5/-5 ORFs) using SILIX and identified their known domains via HMM search (supplementary files S3-5).

We found 644 TpgA homologues widely distributed across *Gamma*- and *Betaproteobacteria* (Figure 2, supplementary Figures S3 to S7). Some homologs contained only a BamE domain, while others had both a BamE domain and a OmpA peptidoglycan-binding domain. SadB (106 homologues) was exclusive to the *Enterobacterales*. Interestingly, in our dataset, genes encoding DUF2827 proteins (327 homologs) were often adjacent to adhesin genes, frequently occuring in two or three successive copies. These DUF2827 proteins, which are rare (around 1000 protein sequences in InterPro Database), are mostly present in *Burkholderia* but also in *Veillonellales* and *Pasteurellales* (Figure 2, supplementary Figure S7). They are cytoplasmic proteins and structural homology points toward a role as a glycosyltransferases (16), suggesting a potential role in TAA glycosylation, though their exact function needs further investigation. Genes encoding the invasion protein B IalB (188 homologs) were commonly linked to TAAs in *Hyphomicrobiales* (supplementary Figure S7). IalB, a small protein associated with adhesion and invasion of human erythrocytes, is located at the inner membrane (17). The presence of IalB often co-occurs with another small protein of unknown function containing a signal peptide. However, IalB is often present without adhesins in its proximity, suggesting that this association may be incidental. Domain analysis did not reveal additional potential candidates (supplementary Table S3) .

**Figure 2:**
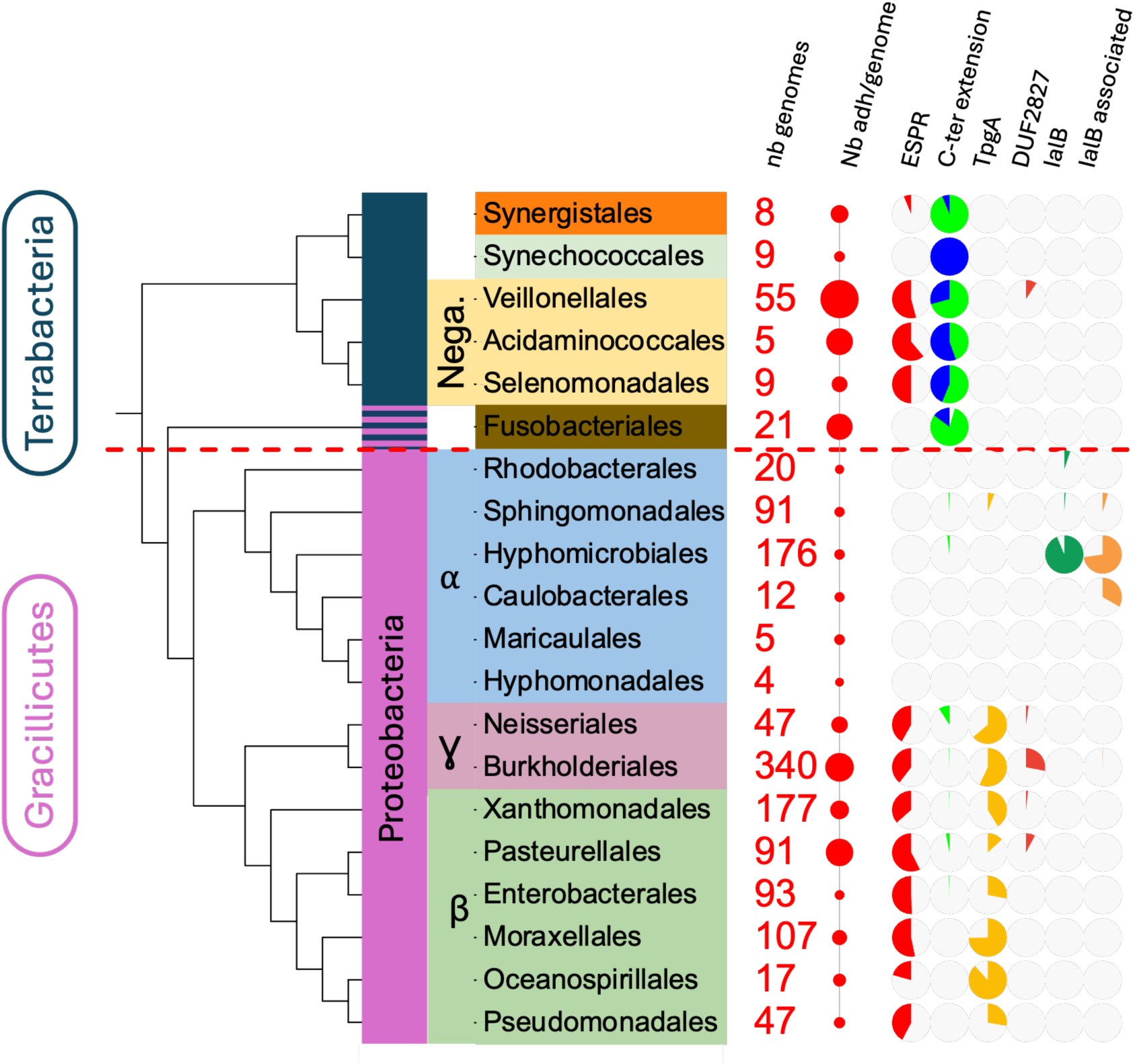
Distribution of the main TAA-associated features. Each piechart represents the percentage of genomes or adhesin containing the corresponding feature. Size of the disc represents the average number of TAA per genome, ranging from 1 for *Hyphomonadales* to 4.3 for *Veillonellales*. For the periplasmic (C-ter) extension, the green and blue colors indicate a short and long periplasmic extensions respectively. The red dotted line represents the separation between Terrabacteria, requiring a PD for their TAAs and Gracillicutes. The assignment of Fusobacteriales to the Terrabacteria remains debated but they share the same PD requirements (37).

### Autotransporters outside of *Proteobacteria* possess periplasmic domains

We wondered if instead of helper proteins, some TAAs could encode for additional undescribed domains helping secretion. Knowing that all TAAs in *V. parvula* possess a periplasmic domain (PD), we first explored the presence and diversity of PDs in trimeric autotransporters across all bacteria by searching TAAs with sequences extending past the C-terminal β-barrel. About 12% (1508) of all TAAs have a C-terminal periplasmic extension of more than 10 residues, almost exclusively outside of Proteobacteria. Nearly all Negativicutes (99.8%) Synergistia (96%) and Synechoccocus (92.5%) and 62% of Fusobacteriia TAAs have an extra PD. In contrast, very few Gammaproteobacteria (74/6655, including 43/406 *Haemophilus*) and Betaproteobacteria (70/2934 including 47/323 *Neisseria*) have a PD (Figures 2 and 3A, supplementary files S4 and S6).

**Figure 3:**
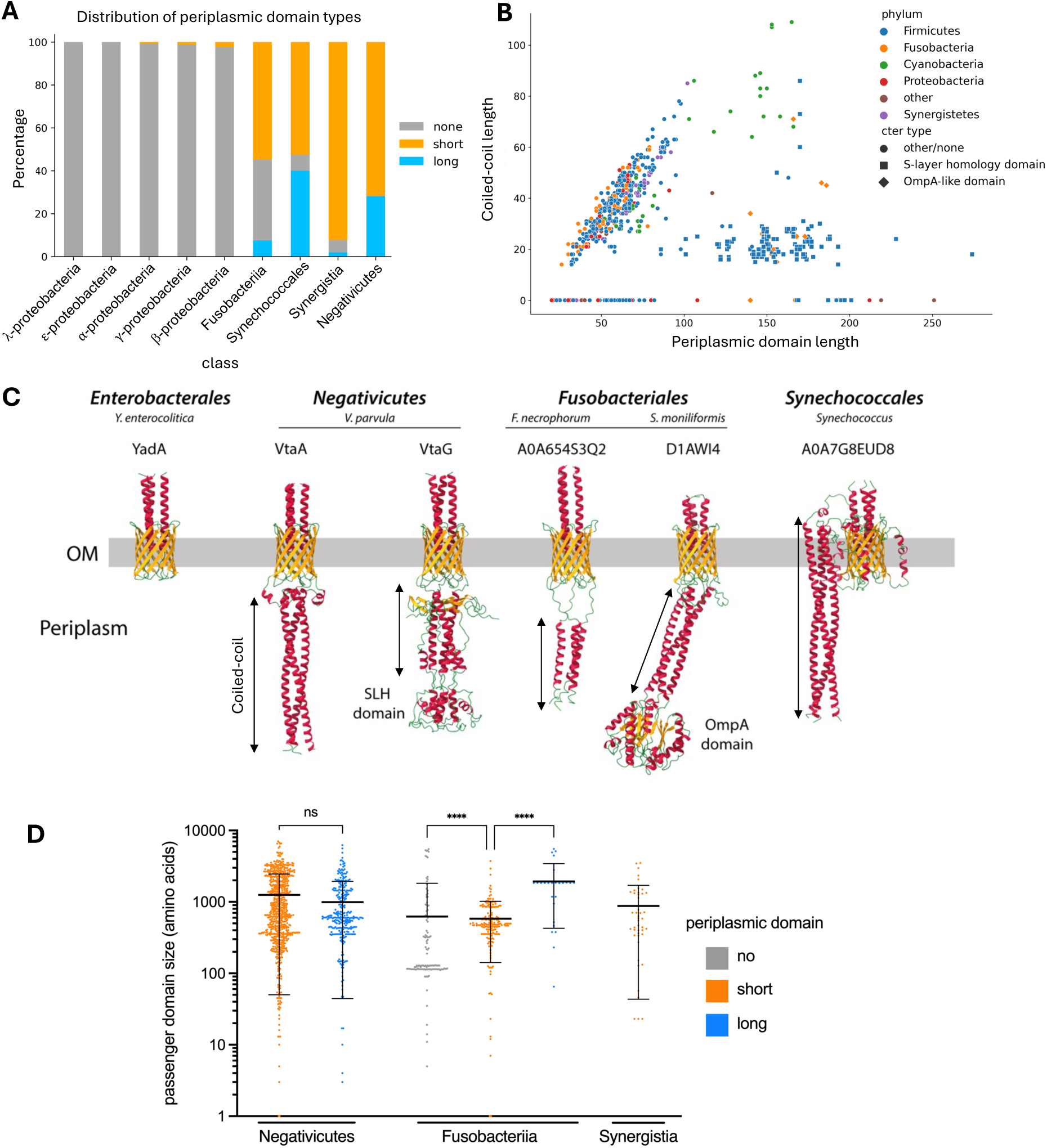
Types of periplasmic domains found in trimeric autotransporters. A) Proportion of the different types of periplasmic domains by class. B) Size of the coiled-coil domains found in the periplasmic domains in function of their size in amino acids. Symbols indicate the additional domains found associated with it and samples are colored by phylum. C) Examples of predicted structure of the periplasmic domains by Alphafold 2.3.2. α-helixes are colored in red, β-sheets in yellow and unstructured regions in green. Black arrows indicate the coiled-coil region of the PDs. The gray bar represents the outer membrane (OM). A0A7G8EUD8 periplasmic domain in *Synechococcales* is mislocalized in the membrane. D) Boxplot representing the size of the adhesin passenger domains, separated by class and type of periplasmic domain. Significance of the differences between adhesins with different PD was calculated using a Kruskal-Wallis test (**** : P ≤ 0.0001).

These PDs come in two distinct types: a “short” version, under 100 residues, mainly consisting of a long coiled-coil domain, and a “long” version, over 100 residues, featuring a short coiled-coil domain followed by a putative peptidoglycan (PG) binding domain. The latter is only found in *Streptobacillus* (Fusobacteriia), where the PG-binding domain is an OmpA domain, and in Negativicutes, where the PG-binding domain is a SLH domain (Figure 3A-C, supplementary Figure S8). The cyanobacteria *Synechococcus* possess long PDs only comprised of a coiled-coil domain (Figure 3B-C). We then investigated if the type of PD domain was associated with a difference in TAA size, similarly to ESPR. In Negativicutes, the type of PD does not appear to correlate with the adhesin size. In Fusobacteriia, adhesins without a PD possess often a very small passenger domain below 100 residues, suggesting inactive adhesins (Figure 3D) and adhesins with a long PD are found only in a few Streptobacillus species, which limits our ability to assess their relationship to overall adhesin size. We could not find any other novel domains of interest in TAAs by HMMer search (see supplementary table S4).

### Periplasmic domains in *V. parvula* are necessary for protein stability

We decided to investigate the function of coiled-coil and SLH PDs using our model strain, the Negativicute *V. parvula* SKV38. We previously identified the functions of three *V. parvula* adhesins containing a coiled-coil domain (supplementary table S5). VtaA mediates autoaggregation (12) and coaggregation with *S. oralis* (18), VtaE mediates coaggregation with *S. gordonii* (18) and VtaF binds to surfaces (Figure 4B). Deleting the C-terminal coiled-coil domain of these adhesins (nucleotides 8935-9123 for *vtaA*, 9253-9426 for *vtaE* and 9445-9579 for *vtaF*) resulted in a loss of function for all (Figure 4A-E), indicating a significant perturbation of the adhesins. We then created PD deletion mutants on adhesins with an additional periplasmic C-terminal HA-tag for detection by Western blot. Without their PDs, the adhesins were either undetectable or present at much lower levels compared to the WT proteins, explaining the observed loss of function (Figure 4F). We also raised an antibody against the VtaA passenger domain (residues 45-700), confirming through Western blot and immunofluorescence that the protein degradation was not due to the presence of the HA-tag (Figure 5A-B). Interestingly, while we did not detect any signal by Western blot targeting the C-terminal HA-tag of VtaA, we observed some degraded protein when detecting the N-terminal passenger domain. It is therefore likely that the protein is degraded first by its C-terminus.

**Figure 4:**
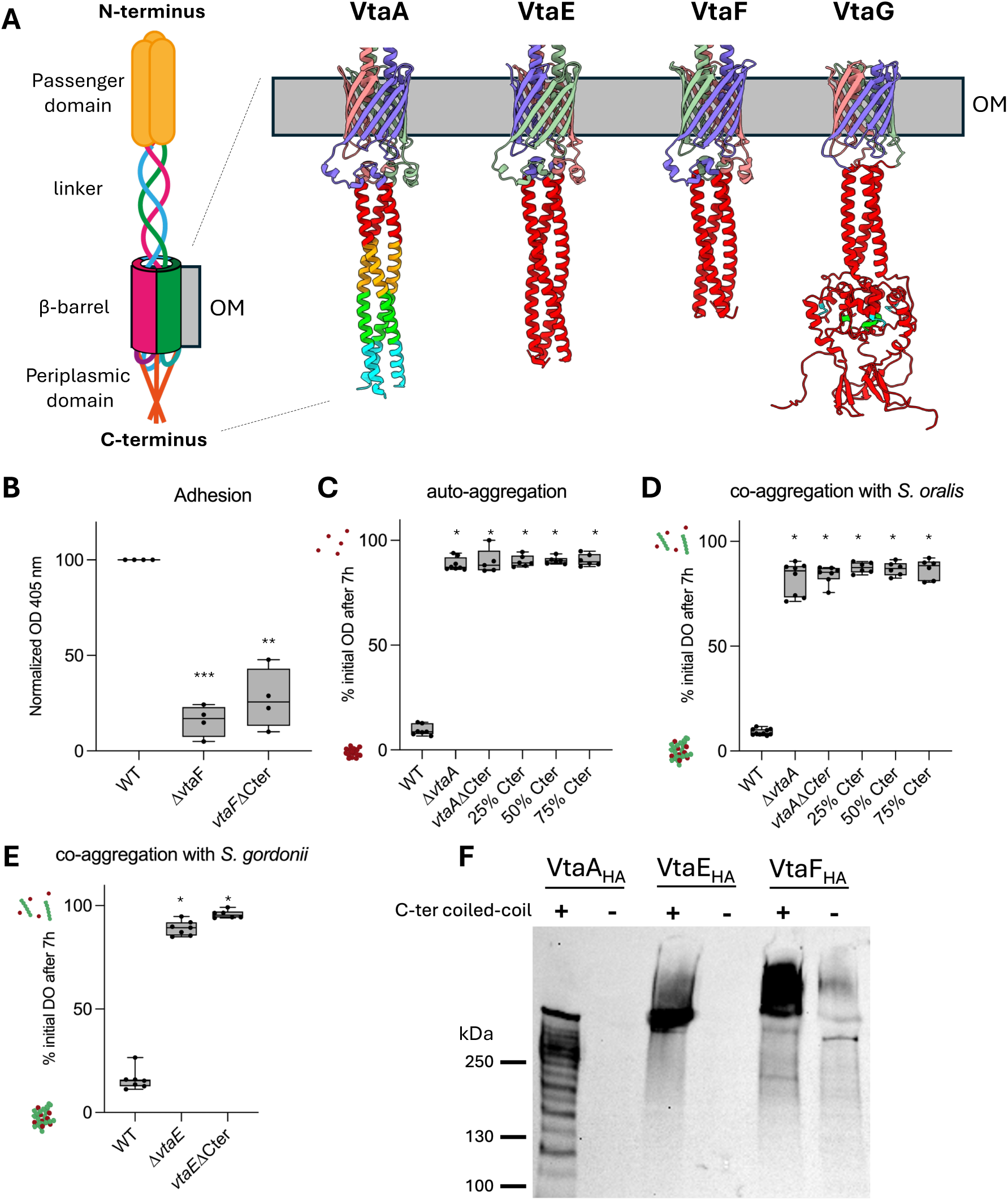
Deletion of the periplasmic coiled-coil domain results in protein instability. A) Schematic representation of *V. parvula* TAA and predicted structure of periplasmic domains by AlphaFold 2.3.2. The deleted section is colored in red. Removal of 25%, 50% and 75% of VtaA coiled-coil correspond to the cyan, green and yellow sections. B) ELISA adhesion assay to Nunc MaxiSorp plates (Invitrogen) precoated with MaxGel ECM mixture (Sigma). The optical density at 405 nm is indicative of the activity of HRP coupled to the antibody and the amount of bacteria adhering to surfaces. All values are normalized to the mean value for WT for each experiment. Experiment performed in 4 biological replicates and technical triplicate or quadruplicate. C-D) Autoaggregation and coaggregation assays with *S. oralis* of the *vtaA* mutants as measured by the decrease in initial optical density after 7h. E) Coaggregation assay with *S. gordonii* of the *vtaE* mutants as measured by the decrease in initial optical density after 7h. For B), stars indicate significatively different conditions compared to the WT, based on a RM one-way ANOVA test corrected for multiple testing ; for C-E) stars indicate significatively different conditions compared to the WT, based on a Kruskal-Wallis test corrected for multiple testing. (*: P ≤ 0.05, **: P ≤ 0.01, ***: P ≤ 0.001). F) Western-blot targeting the HA-tagged adhesins with or without the periplasmic domain using an anti-HA antibody.

**Figure 5:**
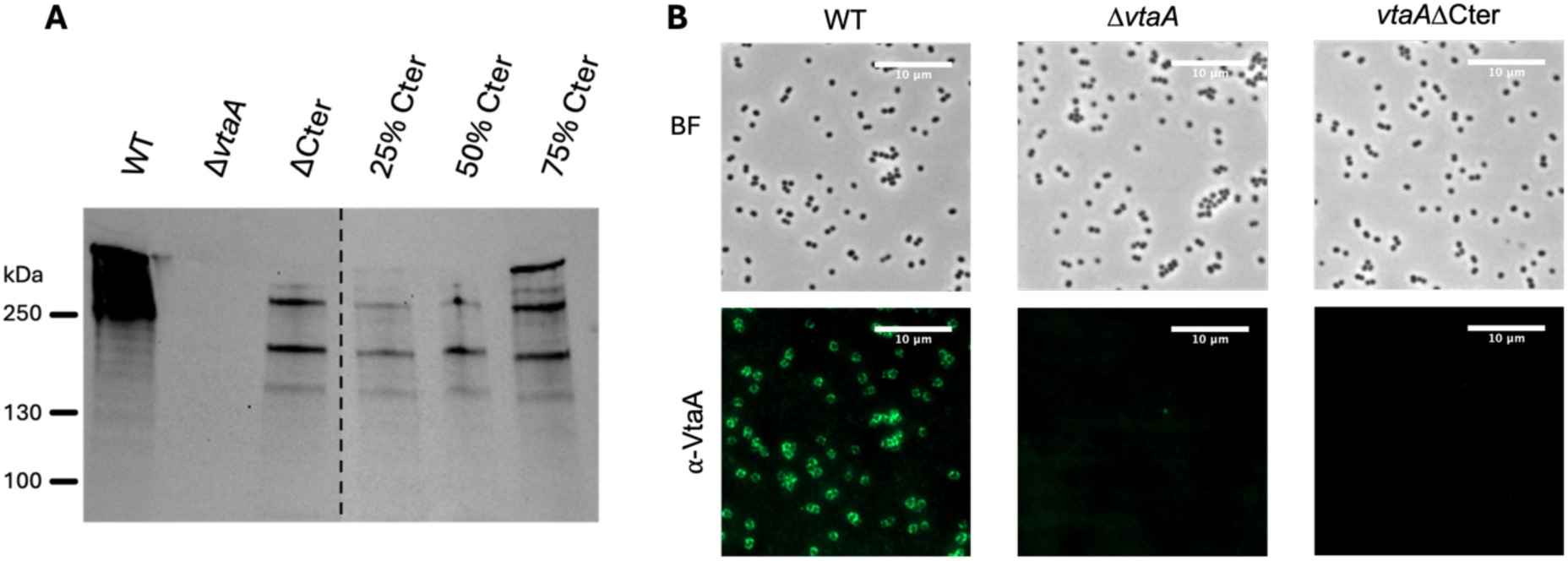
VtaAΔCter is not detected at the cell surface by an antibody targeting its passenger domain. A) Western blot using an anti-VtaA-head antibody on VtaA with or without the periplasmic domain. Membrane was cropped at the dotted line. B) Immunofluorescence using an anti-VtaA antibody on *V. parvula* WT, Δ*vtaA* or *vtaAΔ*Cter. All images were adjusted to the same contrast values using Fiji (38). Scale bar is 10 µm.

Additionally, we performed partial deletions of the coiled-coil domain in VtaA, retaining between 25 and 75% of the domain (Figure 4A, C-D). All modifications resulted in loss of function, an absence of protein on the cell surface and a strong degradation of the protein as observed by Western blot (Figure 5A, supplementary Figure S9).

As an attempt to limit protein degradation in the absence of the PD, we constructed single deletion mutants of four genes coding for putative periplasmic proteases: *bepA* (FNLLGLLA_00052), *rasP* (FNLLGLLA_01074), *hhoB* (FNLLGLLA_00452) and *ctpB* (FNLLGLLA_01133). However, deleting these proteases did not avoid the degradation of VtaA lacking the coiled-coil domain (supplementary Figure S10).

SLH-containing adhesins have no identified function in *V. parvula* SKV38. However, we recently discovered that deleting the putative two-component system FNLLGLLA_00450-51 leads to increased biofilm formation, primarily due to the overexpression of the SLH-containing adhesin VtaG (Audrain et al. in prep). In this Δ*FNLLGLLA_00450-51* background, deletion of the whole *vtaG* gene reduced biofilm formation by ca 50%, while the deletion of the SLH domain of VtaG (*vtaG*ΔSLH, *vtaG* deleted for nucleotides 1240-1731, see Figure 4A) only reduced biofilm formation by 25% (Figure 6A). This indicates that VtaG deleted of its SLH domain is still functional. We constructed strains with a C-terminal HA-tag or with a N-terminal HA-tag right after the signal peptide of VtaG, enabling protein detection at the cell surface. Deletion of the SLH domain resulted in a decreased amount of protein detected by Western blot for both the C-ter or N-ter tagged VtaG (Figure 6B, supplementary Figure S11), consistent with the reduced biofilm formation. VtaG is much smaller (576 amino acids) than VtaA, VtaE and VtaF (each around 3000 amino acids), facilitating detection of trimers on Western blot. VtaGΔSLH still formed trimers on Western blot and was detectable at the cell surface by immunofluorescence, albeit in reduced quantities compared to the WT strain (Figure 6C). Thus, although PDs are not essential for TAA secretion or trimerization in *Veillonella*, they may contribute to these processes by promoting protein stability.

**Figure 6:**
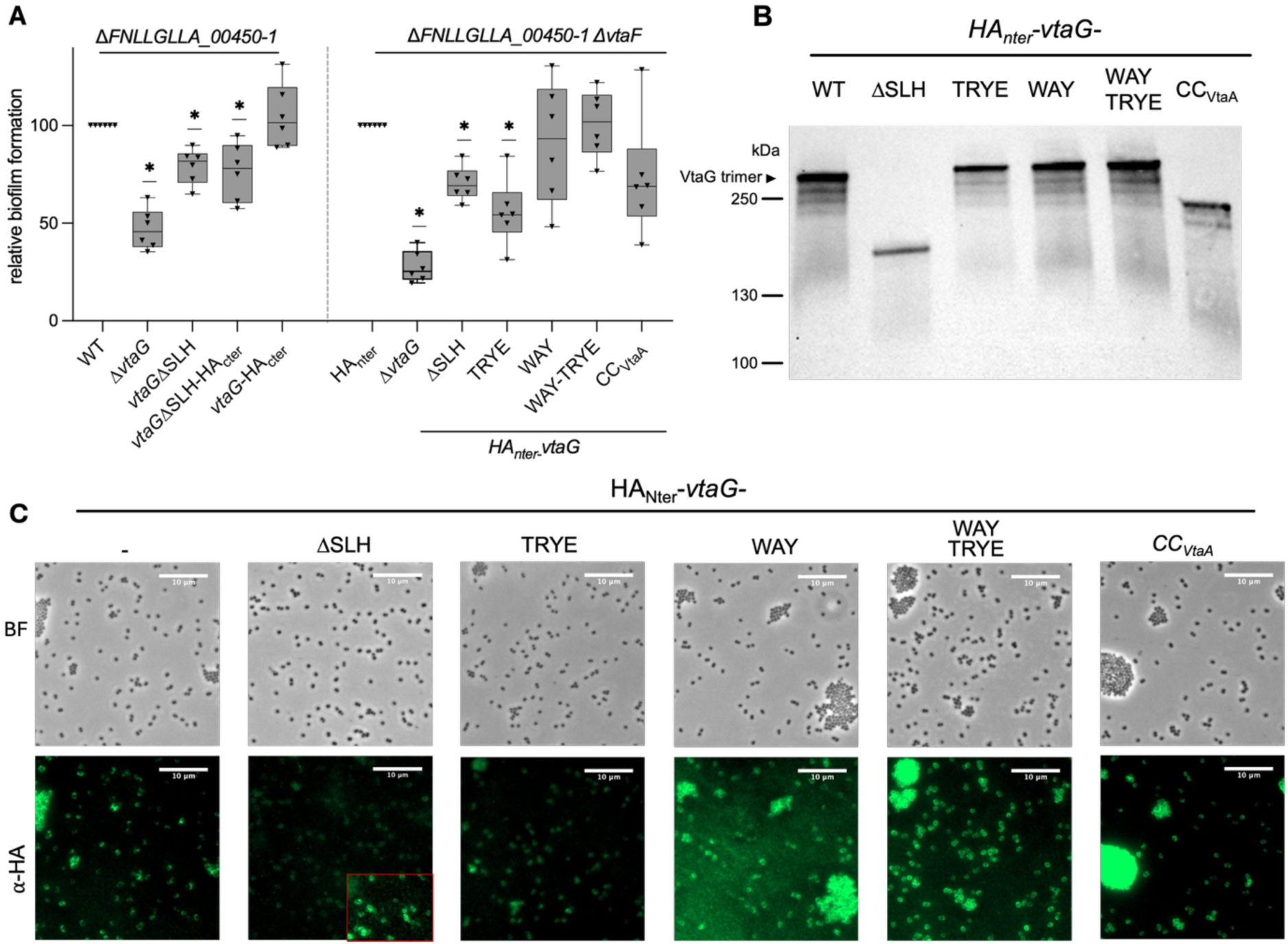
Absence of the periplasmic SLH does not prevent trimerization or secretion. A) Results of a 96 well plate biofilm assay by cristal violet staining after 24h of growth (6 biological replicates). For each replicate, biofilm formation values were normalized to the Δ*FNLLGLLA_00450-51* mutants. * indicates statistical differences (p<0.05) compared to the WT using a Wilcoxon signed rank test. B-C) Western-blot and immunofluorescence using an anti-HA antibody on an HA_Nter_-tagged *vtaG* with various modifications of its SLH domain or *vtaA* coiled-coil inplace of its SLH in a *ΔFNLLGLLA_00450-51::catPΔvtaF::kanR* background. The black arrow in the Western blot indicates the expected trimeric size of VtaG. In immunofluorescence, all images were adjusted to the same contrast values using Fiji (38); for the ΔSLH strain, the red insert correspond to the same image but with increased brightness for better visualization.

### *V. parvula* adhesin SLH domains drive polar localization during division

SLH domains are recognized for their ability to bind peptidoglycan. In *V. parvula* the outer membrane is linked to peptidoglycan modified with polyamines such as cadaverine and putrescine through a trimer of the OmpM porins/OM tether proteins (19,20). These proteins each contain an SLH domain, akin to the SLH-containing TAAs. This led us to investigate whether the SLH domain of TAAs could also bind peptidoglycan. To explore this, we generated mutants of *vtaG*_HAnter_ and *vtaG*_HActer_ with two conserved motifs in the SLH replaced by alanines, WAY_467_AAA and TRYE_496_AAAA (supplementary Figure S12) in a Δ*FNLLGLLA_00450-51 ΔvtaF* background (as *FNLLGLLA_00450-51* tends to form clumps reduced in absence of *vtaF*). Interestingly, the point mutations did not lead to protein degradation, unlike the deletion of the SLH domain (Figure 6B). However, these mutants displayed contrasted phenotypes: the TRYE mutation reduced biofilm formation, whereas the WAY and the double WAY-TRYE mutation did not (Figure 6A). We noticed that the surface-exposed VtaG seemed excluded from the septum during cell division, with numerous individual cells presenting VtaG only on one of their side (Figure 7A). However, all mutations affecting the peptidoglycan binding and the substitution of the SLH domain by VtaA coiled-coil domain resulted in a loss of this polar localization suggesting that PG binding is directly linked to this localization (Figure 7B-D, supplementary Figure S13, supplementary text 1).

**Figure 7:**
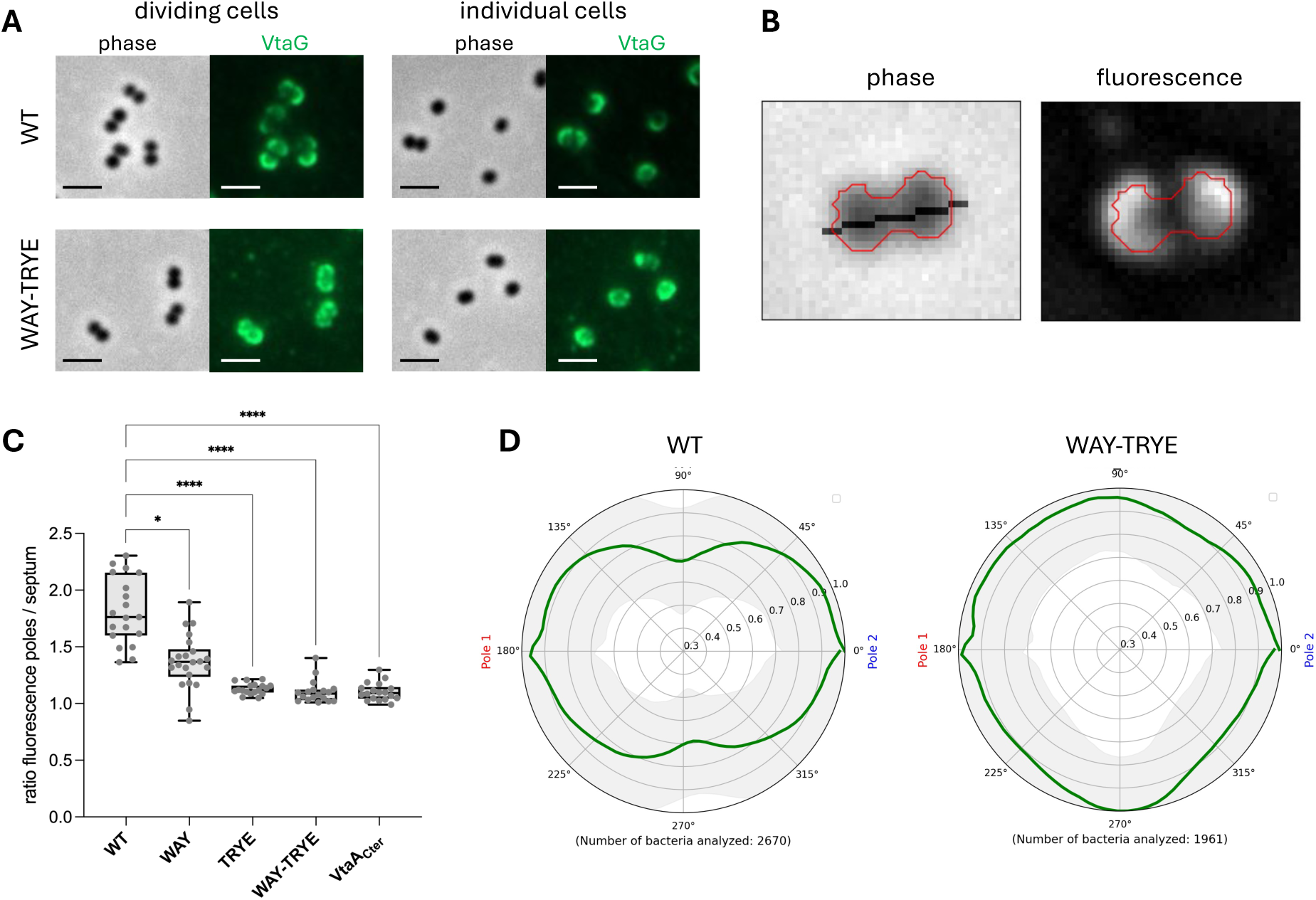
TAAs with an SLH domain are excluded from the septum during division. Immunofluorescence images using an anti-HA antibody on an HA_Nter_-tagged *vtaG* with various modifications of its SLH domain in a *ΔFNLLGLLA_00450-51::catPΔvtaF::kanR* background were analysed to measure septum exclusion of VtaG during division. A) Illustrative images of VtaG localization with either the WT periplasmic domain or the dual WAY-TRYE mutant. Scale bar is 2 μm. All images contrasts were adjusted using Fiji (38). B) Example of the analysis performed on each segmented cell, for which the perimeter and the longer axis (allowing to identify the two poles) was determined on the phase-contrast channel, allowing to retrieve the corresponding fluorescence in the fluorescence channel. C) Average ratio between the normalized fluorescence signal on the poles and the septum calculated with the following formula: 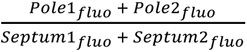 only taking in account dividing cells (defined as having an area above 130 pixels). Each dot is the average for one field of view, with at least 15 replicates per condition on three different biological replicates. Stars indicate significatively different conditions compared to the WT, based on a Kruskal-Wallis test corrected for multiple testing. D) Polar representation of the average relative normalized fluorescence (green line) and its standard deviation (in gray) for the WT periplasmic domain and the dual WAY-TRYE mutant. The septum is approximately located at the 90°-270° axis.

TpgA has been proposed to anchor TAAs to peptidoglycan and facilitate their retention in the cell envelope (9). If the peptidoglycan-binding domain of VtaG serves a similar role, we would expect VtaG to be released in high amount in the extracellular medium when its peptidoglycan binding ability is impaired. However, subjecting *V. parvula* cultures to shear stress by vortexing for 1 minute did not result in increased release of the adhesin into the supernatant for any of the PG-binding defective mutants (Figure 8). This suggests that the SLH domain of VtaG does not significantly contribute to its anchoring in the cell envelope.

**Figure 8:**
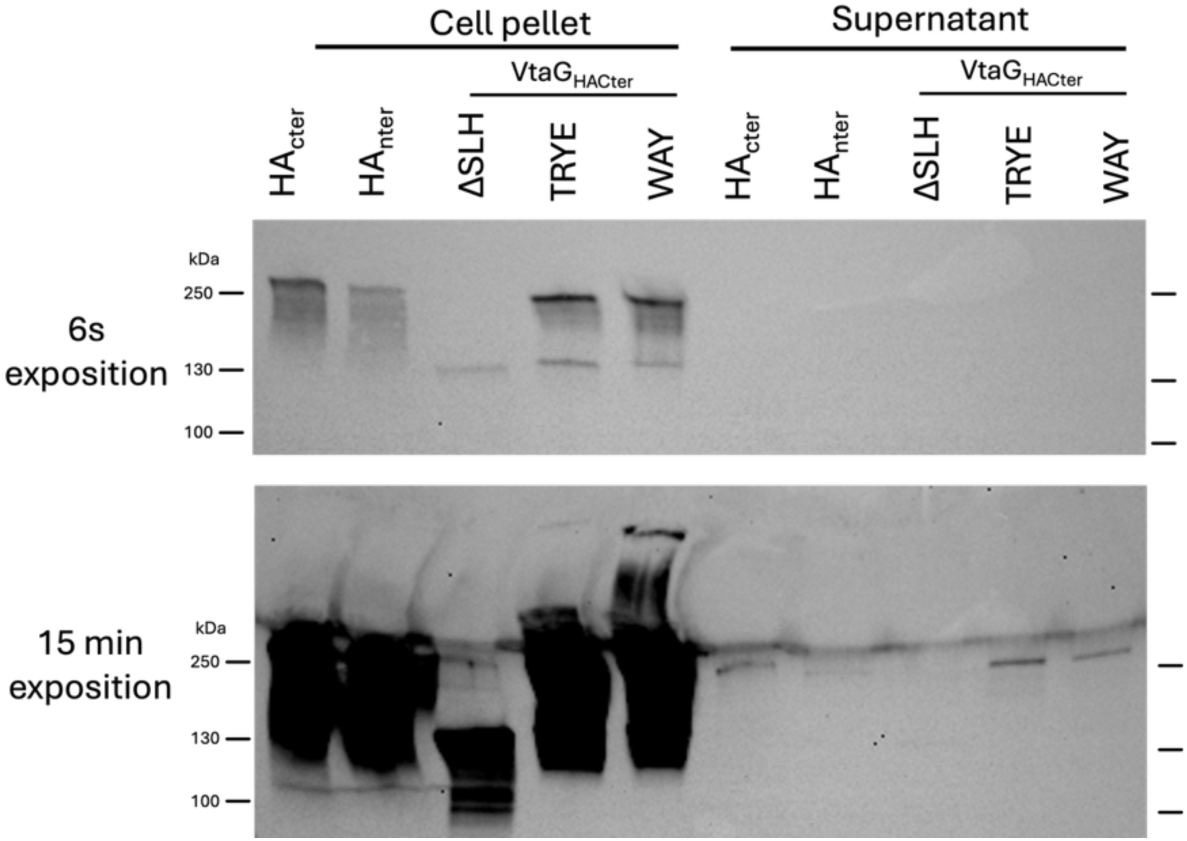
the SLH domain does not seem to favor anchoring of VtaG to the cell envelope. Western blot using an antibody targeting the HA-tag present in C-terminus or N-terminus of VtaG with different SLH modifications. Each sample was separated between the cell pellet and the corresponding TCA precipitated supernatant. The same membrane was exposed with different exposure times (6s or 15 min).

### Periplasmic domains are associated to adhesin function

We investigated the contribution of PDs to adhesin function by substituting the coiled-coil domain of VtaA with the one of VtaF and two SLH domains from VtaH and FNLLGLLA_00037, a partial adhesin probably unable to reach the periplasm because of absence of signal peptide (12) (Figure 9A). Introducing an SLH domain to VtaA markedly impaired auto-aggregation, resembling phenotype of a Δ*vtaA* strain (Figure 9A). We have previously constatated that auto-aggregation most likely requires more VtaA at the cell surface than co-aggregation, which is less sensitive to a decrease of the quantity of VtaA (12,18). Indeed, the same mutants presented only a slight reduction in co-aggregation compared to the WT (Figure 9B), suggesting a reduced quantity of the hybrid VtaA compared to the WT version. In contrast, replacing the longer coiled-coil domain of VtaA with that of VtaF impacted much less VtaA activity with only a partial reduction in auto-aggregation and no effect on co-aggregation (Figure 9A-B). Immunofluorescence and Western blot analyses indicated that substituting the PD both by an SLH domain or another coiled-coil domain resulted in lower production and surface exposure of proteins (albeit less extensively for the coil-coiled domain), potentially explaining the reduced auto-aggregation (Figure 9D-E). We conducted a similar experiment by replacing the SLH domain of VtaG with the coiled-coil domain of VtaA. This modification did not lead to a significant decrease in biofilm formation, although there was possibly a slight protein degradation observed in Western Blot (Figure 6A-B). More importantly, VtaG subsequently loose its specific localization, suggesting that TAAs periplasmic coiled-coil domains are unable to bind to PG (Figure 6). In conclusion, it appears that PDs co-evolved with their cognate beta-barrels and are not always interexchangeable.

**Figure 9:**
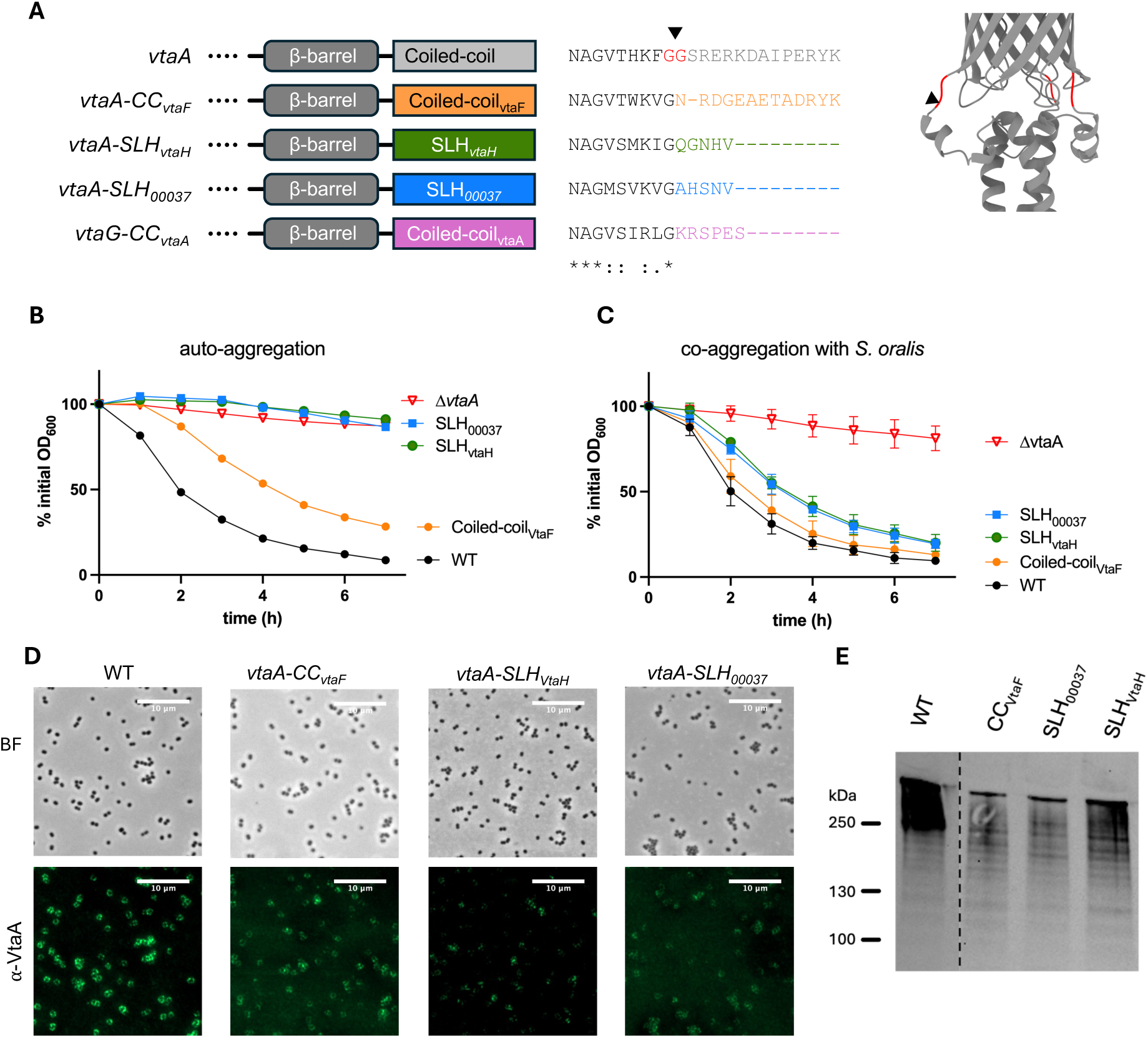
Periplasmic domains are not totally interchangeable between adhesins. A) Schematic representations of the different constructed chimeric adhesins. The exact region exchanged is indicated by a black arrow on an alignment of the four adhesin sequences and on the model of VtaA β-barrel. B-C) auto-aggregation and co-aggregation assay for the different hybrid constructs. Experiment done in 3-8 independent biological replicates, error bars represent the standard deviation. D-E) Immunofluorescence images and Western blot using an anti-VtaA antibody on the different mutants. The WT and different mutants were on the same membrane that was cropped (same membrane used in Figure 5a).

## Discussion

Adhesion plays a crucial role in bacterial biology, facilitating biofilm formation and enhancing virulence. Autotransporters, including trimeric ones, are key surface-exposed adhesins often requiring , in addition to their standard anchor, stalks and passenger domains, additional helper proteins and domains. Here, we described additional PDs found in nearly all TAAs outside of *Proteobacteria*, some of which potentially binding to peptidoglycan.

In *V. parvula*, deleting the PDs of TAAs leads to protein instability and degradation. This finding is puzzling because *Proteobacteria* TAAs, which are structurally similar to *V. parvula* TAAs, do not rely on PDs for stability, suggesting that PD likely fulfill other functions than only protein stability. Overall, our bioinformatic analysis highlights a clear divide between Proteobacteria and Terrabacteria, where TAAs require a PD. This suggests that these PDs might compensate for the absence of certain elements present in Proteobacteria’s membrane systems. The major membrane system important for TAAs secretion is the BAM machinery. Negativicutes, Cyanobacteria and Fusobacteriia lack the full complement of lipoproteins found in Proteobacteria’s BAM machinery, relying solely on the core BamA (21). However, BamD, conserved in all Gracilicutes, has been described in *E. coli* as essential for the secretion of trimeric autotransporters (22) and the widely distribute helper protein TpgA possess a BamE-like domain with an additional OmpA peptidoglycan-binding domain (8). It is plausible that TAAs PDs facilitate interactions with BamA in Terrabacteria and accelerate secretion through mechanisms that remain to be elucidated.

We showed that TAAs deleted from their PDs, at least for the SLH domain, still trimerize and are secreted, indicating that the PD is not absolutely necessary for these processes. However, these domains might assist or accelerate trimerization and subsequent secretion. Their absence could lead to partially unfolded adhesins in the periplasm, resulting in enhanced degradation. According to the accepted TAA secretion threading model (23), the unfolded passenger domain translocates through the β-barrel lumen emerging in the periplasm. This means that unfolded passenger domain would locate below the β-barrel lumen, around where the PDs are predicted to trimerize. Thus, these PDs may remain physically hindered from trimerizing until the passenger domain is fully secreted. It remains unclear whether these PDs facilitate protein secretion or only become functionally important only once the adhesin is properly assembled.

Inverted autotransporters have two types of PDs: a regular alpha helix of unknown function or a LysM domain facilitating peptidoglycan binding, surface exposure and dimerization (10,11). TAAs also present this dichotomy with either a PD made of three alpha-helixes or a PG-binding domain. AlphaFold modeling suggests that both types of TAAs PDs may trimerize by forming a superhelix made of three alpha helixes. The size, sequence, and final residue of the short PD are poorly conserved, even within *Veillonella* species (supplementary Figure S12), which points to a structural function. The non-specific localization of VtaG fused to the VtaA coiled-coil domain additionnaly supports the idea that short PDs do not bind peptidoglycan. To study the PG-binding domain function, we both deleted the full SLH domain and did point mutations inactivating the PG-binding capacity. While the former resulted in strong protein degradation, suggesting a role for stability, the latter only affected the TAA polar localisation during division. These results point towards a bifunctional role for PG-binding PDs, one of stability (possibly due to the trimerization of the alpha-helix), and the other of PG anchoring. The septum exclusion of PG-binding TAAs is a puzzling result, both by its function, and the underlying mechanism. One first hypothesis to explain the PG-binding dependent exclusion of the septum during division is that PG binding prevents adhesins to laterally diffuse to the nascent OM. OMPs are largely immobile in *E. coli* (24), but *V. parvula* may have an increased membrane fluidity allowing OMP diffusion as described in *M. xanthus* (25). SLH domains could possibly also have a higher affinity for mature peptidoglycan, which would favor their periplasmic localization and subsequent incorporation by BamA outside the poles. Intriguly, in *E. coli*, BamA is thought to be more active at the septum (26), which might not be the case in *Veillonella*.

TAAs PDs are primarily found in adhesins outside of Proteobacteria, with peptidoglycan binding capacity emerging twice independently in Fusobacteriia (with the OmpA domain) and in Negativicutes (with the SLH domain), which reflects their respective main outer membrane tethering systems (19). Unlike Proteobacteria, Terrabacteria rely exclusively on OmpM, a trimeric porin associated with an SLH domain (19–21), for membrane tethering, while Fusobacteriia use OmpA domains. The PG-binding domain of TAAs may serve as an additional anchoring mechanism to prevent outer membrane tearing by shear stress exerted by the adhesin, a hypothesis previously suggested for the TpgA helper protein (9). However, their potential role in membrane attachment seems limited, as deletion of *ompM* genes alone in *V. parvula* is sufficient to induce an almost complete membrane detachment (19).

In our analysis of adhesin-containing genomes, we did not identify new proteins associated with the secretion of trimeric autotransporters. While we found SadB and TpgA homologues in many genomes, no other potential candidates emerged. Most identified genes were linked to regulation and signal transduction and were not specifically adjacent to adhesin encoding genes (supplementary Table S3). The most promising candidate are proteins of the DUF2827 family, whose function remains unknown, but that are frequenctly associated with TAA adhesins in *Burkholderia* but also in rare cases in *Veillonellales* and *Pasteurellales*; however, their cytoplasmic localization suggests that they are not involved in secretion through the OM but could be important for post translational glycosylation modifications. It is likely that the genes encoding these DUF2827 proteins have been horizontally acquired in *Veillonella*, probably independently of their neighbouring TAA encoding genes regarding the low homology between TAA adhesins in *Burkholderia* and *Veillonella* and the presence of a PD in *Veillonella* TAAs found next to DUF2827 proteins. It seems unlikely that additional helper proteins await discovery, prompting questions about why only certain TAAs require them.

A few coiled-coil PDs are found in Proteobacteria. It would be interesting to trace their origins to determine if those are the result of an horizontal transfer or an independent convergent evolution of the TAAs. However, our attempt to reconstruct the phylogeny of β-barrels with PDs, based on their C-terminal sequences, including the β-barrel and PD sequences, was unsuccessfull (supplementary Figure S14). The phylogeny was poorly supported by bootstrap analysis, and there was no clear indication of the origin of the *Proteobacteria*l PDs. Additionnaly, we could not also establish a clear relationship between the different PDs found in other bacteria. Overall, we were not able to establish the origin of the common ancestor of trimeric autotransporters and whether it already possessed PDs or not.

The ESPR is believed to limit the rate of secretion through the SEC translocon to reduce the risk of misfolding (13). Consistently, we found that the presence of the ESPR correlates with longer adhesins on average both for monomeric and trimeric autotransporters. We determined that both Alphaproteobacteria and Fusobacteriia lack the ESPR signal in their trimeric and classical autotransporters, consistent with prior findings for Alphaproteobacteria (15). However, Negativicutes possess ESPR in both classes. Interestingly, Negativicutes are phylogenetically more distant from Gamma and Beta Proteobacteria than Fusobacteriia and Alphaproteobacteria. Investigating the evolution of the SEC machinery in these diverse bacterial groups could provide valuable insights on their differences in mechanisms of action.

In conclusion, the “universal” secretion machinery of TAAs is clearly not so general and has adapted differently to its different bacterial host. TAAs in Terrabacteria have consistently developed PDs, essential for protein stability but sometimes also involved in peptidoglycan binding. On the other hand, the ESPR domain has not been coopted by Fusobacteria and Alphaproteobacteria. These variations in TAA secretion mechanism might reflect the underlying differences in the secretion machineries (SEC and BAM machineries) of their different host. Future work will be required to investigate the interplay between additional domains of TAAs and the different secretion machineries and complement missing components in different bacteria.

## Methods

### Growth conditions

Bacterial strains are listed in table S1. *Streptococcus spp.* were grown in brain heart infusion (BHI) medium (Bacto brain heart infusion; Difco). *V. parvula* was grown in SK medium (10 g liter^−1^ tryptone [Difco], 10 g liter^−1^ yeast extract [Difco], 0.4 g liter^−1^ disodium phosphate, 2 g liter^−1^ sodium chloride, and 10 ml liter^−1^ 60% [wt/vol] sodium DL-lactate; described in Knapp et al.(27)) or in BHI supplemented with 0.6% sodium DL-lactate (BHIL) when observing autoaggregative phenotypes. Bacteria were incubated at 37°C under anaerobic conditions in anaerobic bags (GENbag anaero; bioMérieux no. 45534) or in a C400M Ruskinn anaerobic-microaerophilic station. *Escherichia coli* was grown in lysogeny broth (LB) (Corning) medium under aerobic conditions at 37°C. When needed, 20 mg/L chloramphenicol (Cm), 200 mg/L erythromycin (Ery), 300 mg kanamycin (Kan) or 2.5 mg/L tetracycline (Tet) was added to *V. parvula* cultures, 25 mg/L Cm or 100 mg/L ampicilin (Amp) was added to *E. coli* cultures. All chemicals were purchased from Sigma-Aldrich unless stated otherwise.

### *Veillonella parvula* natural transformation

From plate, cells were resuspended in 1 ml SK medium adjusted to an optical density at 600 nm (OD_600_) of 0.4 to 0.8, and 15 μl was spotted on SK agar petri dishes. On each drop, 1-5 μl (75 to 200 ng) linear double-stranded DNA PCR product was added. The plates were then incubated anaerobically for 24-48 h. The biomass was resuspended in 500 μL SK medium, plated on SK agar supplemented with the corresponding antibiotic, and incubated for another 48 h. Colonies were streaked on fresh selective plates, and the correct integration of the construct was confirmed by PCR and sequencing.

### *Veillonella parvula* mutagenesis and complementation

*V. parvula* site directed mutagenesis was performed as described by Knapp and al (27) and Béchon et al (12). Briefly, upstream and downstream homology regions of the target sequence and the *V. atypica* kanamycin (*aphA3* derived from the pTCV-erm(28) plasmid under the *V. parvula* PK1910 *gyrA* promoter) or tetracycline resistance cassette were PCR amplified with overlapping primers using Phusion Flash high-fidelity PCR master mix (Thermo Scientific, F548). PCR products were used as templates in a second PCR round using only the external primers, resulting in a linear dsDNA with the antibiotic resistance cassette flanked by the upstream and downstream sequences. For insertion of the HA-tags, primers containing the HA-tag sequence in their overhangs were used to construct the upstream homology region. For construction of VtaG-HA_nter_, the full *vtaG-HA_nter_* gene was inserted in a Δ*vtaG::eryR* strain to prevent recombination with the C-terminus region of the adhesin only. Primers used in this study are listed in Table S2 in the supplemental material.

### Aggregation assay

Overnight cultures were centrifuged for 5 min, 5000 g and resuspended in aggregation buffer(29) (1 mM Tris-HCl buffer, pH 8.0, 0.1 mM CaCl_2_, 0.1 mM MgCl_2_, 150 mM NaCl) to a final OD_600_ of 1. 400 μL of each culture for coaggregation or 800 μL for auto-aggregation were added to a microspectrophotometer cuvette (Fisherbrand) and left to sediment on the bench in the presence of oxygen, so no growth should occur. The OD_600_ was measured every hour in a single point of the cuvette using a SmartSpec spectrophotometer (Bio-Rad). OD_600_ were then normalized to the initial OD_600_.

### Purification of *vtaA* head domain and antibody generation

Nucleotides 133 to 2100 (5’, outside the ESPR signal sequence) of *vtaA* were amplified by PCR and then integrated by Gibson reaction into the plasmid pET22b, under the control of a T7 promoter regulated by the lac operator. The plasmid was transformed by electroporation into the *E. coli* strain DH5α. After selection for transformation using ampicillin, the presence of the construct in the plasmid was verified by PCR and sequencing. The plasmid purified by Miniprep (Qiagen) was transformed into *E. coli* BL21DE3 pDIA17, the latter carrying l*acIq*. One colony was used to inoculate an overnight culture of LB + Amp + Cm. 100 mL of antibiotic-supplemented culture was inoculated at OD 0.05. Protein production was induced by adding 0.1 mM IPTG when the culture reached OD 0.2. After 3 h of growth at 30°C, the culture was harvested and the protein purified using HisLink^TM^ Protein Purification Systems (Promega) resin columns before dialysis against a 150 mM NaCl, 20 mM Tris-HCl buffer. The purified proteins were assessed for purity via SDS-PAGE and were quantified with the Bicinchoninic Acid Protein Assay Kit (Sigma). Rabbit polyclonal antisera were raised against the recombinant protein using four immunizations (CovalAb). The antisera were adsorbed twice against a crude protein extract of *V. parvula* Δ*vtaA* before use.

### Surface attachment

Bacterial attachment was tested by ELISA in microtiter plates. Plates (MaxiSorp , Nunc) were covered overnight at 4°C with 150 μL of MaxGel Human ECM (Sigma-Aldrich) diluted 20-fold in carbonate buffer (pH 9, 50 mM). The wells were washed three times with 300 μL of PBS then blocked with PBS-1% milk for 1 h. After three washes with PBS, 100 μL of bacterial cell suspension (normalised to OD600 = 0.1) in PBS-0.2% milk was added and the plates were incubated at 37°C for 2 hr. After 2 washes with PBS to remove non-adherent bacteria, the remaining cells were fixed with 4% paraformaldehyde (PFA) in PBS for 30 min at RT, washed and incubated for 1 h with 100 μL of anti-*V. parvula* antibody diluted to 250 ng/mL in PBST-0.2% BSA. After three washes with PBST, cells were incubated for 1 h with an HRP-coupled anti-rabbit secondary antibody (Cell signaling technology, 7074S) diluted 1:1000 in TBST-130.2% BSA. After three final washes in PBST, adherent bacteria were detected by adding 150 μL of ABTS solution (Sigma A3219), and absorbance was read at 405 nm.

### Immunofluorescence and microscopy protocol

30 μL of bacterial culture was deposited on each well of a slide previously covered with L-poly-lysine, before incubation for 5 min at room temperature. 15 μL were removed before being replaced with 4% PFA, 2 times. After removal of all the PFA, the wells were covered with PFA and incubated for 10 min at room temperature. The PFA was then removed and replaced with 50 mM NH_4_Cl in PBS and incubated for 3 min at room temperature, followed by three washes with PBS. For immunofluorescence using anti-VtaA antibodies, the wells were then coated with 0.5% PBS-BSA for 15 min at room temperature, before adding the anti-VtaA antibody (diluted 1:200) in 0.5% PBS-BSA to the wells and incubating for 45 min at room temperature. After three washes with PBS, a secondary anti-rabbit antibody coupled with the fluorophore Alexa488 (Invitrogen, A11034) diluted at 1/300 in 0.5% PBS-BSA was added before incubation for 45 min at room temperature in the dark. After three washes with PBS and one wash with water, 3 μL of Dako fluorescent mounting media (Agilent) was added to each well. For immunofluorescence targeting of HA labels, an anti-HA antibody (Novus Biologicla, NB-600-363) diluted 1:250 instead of the anti-VtaA antibody was used. Cells were imaged using a Zeiss Axioplan 2 microscope equipped with an Axiocam 503 mono camera (Carl Zeiss, Germany). Epifluorescence images were acquired using the ZEN lite software (Carl Zeiss, Germany) and processed using Fiji (ImageJ).

### Protein detection by western blot

Cultures were grown overnight in SK medium. For each culture, an equivalent of 1 mL at OD600 of 2 was centrifuged (5000 g for 5 min), resuspended in 100 μL of Laemmli buffer and incubated for 5 min at 98°C. When needed, supernatants were filtered using 0.2 µm filters, precipitated by addition of 10% TCA final and incubation on ice for one hour. The supernatants were then centrifuged (15 000 g for 10 min), washed with acetone, centrifuged again and resuspended in Laemmli buffer. 10 μL of these cell extracts were added on 12 Mini-Protean TGX stain-free precast gel (Bio-Rad) in 1X TGX buffer (migration: 170 V, 45 min), then transferred on a nitrocellulose membrane using a PowerBlotter XL system (ThermoFisher). For blocking, the membrane was incubated for 1 h with PBST (0.05% Tween - 1 X PBS) with 5% milk. The membrane was incubated for one hour with an anti-HA (NB-600-363, Novus) or an anti-vtaA antibody diluted 2500-fold and 1000-fold respectively in PBST-1% milk. After a second blocking step, the membrane was incubated with a coupled anti-IgG_rabbit_-HRP antibody. After 4 rinses in PSBT, proteins were visualized using ECL^TM^ Western Blotting Detection Reagents (Amersham^TM^).

### Bioinformatic analysis of the periplasmic domains

All proteins sequences possessing a YadA-anchor domain (IPR005594) were downloaded from the InterPro database (https://www.ebi.ac.uk/interpro/entry/InterPro/IPR005594/) as long as all their associated domains (Table S3). The resulting information was analyzed using personalized python scripts. All sequences found in C-terminus downstream of the YadA-anchor domain and with a length above 5 residues were considered as periplasmic domains. Phylogenetic information was extracted using the Pytaxonkit library (30). Coiled-coil predictions was done using DeepCoil2 locally (31) on the periplasmic domain sequences. Structure predictions were done using AlphaFold-Multimer 2.3.2 locally (32) and visualized using USCF ChimeraX 1.6 (33) and PyMOL 2.4.2 (Schrodinger, LCC).

### Bioinformatic analysis of the neighboring genes

We selected all adhesins possessing both a YadA-anchor and a YadA-head domain (IPR008640) in the InterPro database (7803 sequences) and retrieved the NCBI identifier corresponding to their genome of origin, resulting in 2160 genomes. We subsequently filtered them by removing genomes excluded from RefSeq and those with adhesins presenting identical sequences on the 100 last residues before the end of the β-barrel, resulting in 1384 genomes. Adhesin neighboring genes (five upstream and five downstream) were extracted from genomes and their domains analyzed by HMMscan locally with defaults settings. Additionally the neighboring genes were clustered using SiLiX (34) using default parameters. To build the bacterial phylogeny, we divided the genomes among *Gammaproteobacteria* (504 genomes), *Betaproteobacteria* (330 genomes), *Alphaproteobacteria* (281 genomes) and the rest (101 genomes). HMM-based homology searches (with the option --cut_ga) were carried out with HMMSEARCH using the Pfam profiles PF04997.12, PF04998.17, PF04563.15, PF00562.28 and PF11987.8 corresponding to RNA polymerase subunits β, β′ and translation initiation factor IF-2. Single genes were aligned using MAFFT v.7.407 with the L-INS-I option and trimmed using BMGE-1.12 with the BLOSUM30 substitution matrix. The resulting trimmed alignments were concatenated into a supermatrix and the tree built using IQ-TREE (35) with the best-fit model estimated by ModelFinder, and ultrafast bootstrap supports computed on 1,000 replicates of the original dataset. Phylogenetic trees visualization and annotation was done using iTOL 6.9.1 (Minh et al. 2020).

## Data availability

Supplementary datasets used in figure 1-3 and figure 7 are available at: https://github.com/ldorison/periplasmic_domains_TAAs

## Acknowldegments

We thank Najwa Taib and Simonetta Gribaldo for their suggestions for the phylogeny analysis. We thank Nicolas Desprat for his advice on image analysis. This work was supported by Institut Pasteur and grants by the French government’s Investissement d’Avenir Program, Laboratoire d’Excellence “Integrative Biology of Emerging Infectious Diseases” (grant n°ANR-10-LABX-62-IBEID). L.D. was supported by a MENESR (Ministère Français de l’Education Nationale, de l’Enseignement Supérieur et de la Recherche) fellowship.

L.D., C.B. designed the experiments. L.D., S.C.-R. and B.A performed the experiments. L.D. and C.B. wrote the paper, with contributions from B.A and J.-M.G. All authors read and approved the manuscript.

## Notes

https://github.com/ldorison/periplasmic_domains_TAAs

### Competing Interest Statement

The authors have declared no competing interest.

## Bibliography

1. Leo J, Grin I, Linke D. Type V secretion: mechanism(s) of autotransport through the bacterial outer membrane. Philos Trans R Soc Lond B Biol Sci. 2012;367(1592):1088–101.

2. Selkrig J, Mosbahi K, Webb CT, Belousoff MJ, Perry AJ, Wells TJ, et al. Discovery of an archetypal protein transport system in bacterial outer membranes. Nat Struct Mol Biol. 2012 Apr 1;19(5):506–10, S1.

3. Yang B, Fan R, Batista MB, Chen Y, Duan X, Wang R, et al. Structural basis of outer membrane biogenesis by the TamAB translocase. Nat Commun. 2026 Jan 13;17(1):437.

4. Wang X, Nyenhuis SB, Bernstein HD. The translocation assembly module (TAM) catalyzes the assembly of bacterial outer membrane proteins in vitro. Nat Commun. 2024 Aug 23;15:7246.

5. Desvaux M, Scott-Tucker A, Turner SM, Cooper LM, Huber D, Nataro JP. A conserved extended signal peptide region directs posttranslational protein translocation via a novel mechanism. Microbiology. 2007;153(Pt 1):59–70.

6. Cotter SE, Surana NK, St. Geme JW. Trimeric autotransporters: A distinct subfamily of autotransporter proteins. Trends in Microbiology. 2005;13(5):199–205.

7. Grin I, Hartmann MD, Sauer G, Hernandez Alvarez B, Schütz M, Wagner S, et al. A Trimeric Lipoprotein Assists in Trimeric Autotransporter Biogenesis in Enterobacteria. J Biol Chem. 2014 Mar 14;289(11):7388–98.

8. Ishikawa M, Yoshimoto S, Hayashi A, Kanie J, Hori K. Discovery of a novel periplasmic protein that forms a complex with a trimeric autotransporter adhesin and peptidoglycan. Molecular Microbiology. 2016;101(3):394–410.

9. Yoshimoto S, Suzuki A, Kanie J, Koiwai K, Lupas A, Hori K. Insights into the complex formation of a trimeric autotransporter adhesin with a peptidoglycan-binding periplasmic protei [Internet]. 2023 [cited 2024 Jan 10]. Available from: 10.1101/2023.12.21.572085

10. Martinez-Gil M, Goh KGK, Rackaityte E, Sakamoto C, Audrain B, Moriel DG, et al. YeeJ is an inverse autotransporter from Escherichia coli that binds to peptidoglycan and promotes biofilm formation. Scientific Reports. 2017 Sept 12;7(1):11326.

11. Leo JC, Oberhettinger P, Chaubey M, Schütz M, Kühner D, Bertsche U, et al. The Intimin periplasmic domain mediates dimerisation and binding to peptidoglycan. Molecular Microbiology. 2015;95(1):80–100.

12. Béchon N, Jiménez-Fernández A, Witwinowski J, Bierque E, Taib N, Cokelaer T, et al. Autotransporters Drive Biofilm Formation and Autoaggregation in the Diderm Firmicute Veillonella parvula. Journal of Bacteriology [Internet]. 2020 Oct 8 [cited 2021 Feb 23];202(21). Available from: https://jb.asm.org/content/202/21/e00461-20

13. Dautin N. Folding Control in the Path of Type 5 Secretion. Toxins. 2021 May;13(5):341.

14. Szabady RL, Peterson JH, Skillman KM, Bernstein HD. An unusual signal peptide facilitates late steps in the biogenesis of a bacterial autotransporter. Proceedings of the National Academy of Sciences. 2005 Jan 4;102(1):221–6.

15. Desvaux M, Cooper LM, Filenko NA, Scott-Tucker A, Turner SM, Cole JA. The unusual extended signal peptide region of the type V secretion system is phylogenetically restricted. FEMS Microbiol Lett. 2006;264(1):22–30.

16. Campos CG, Byrd MS, Cotter PA. Functional Characterization of Burkholderia pseudomallei Trimeric Autotransporters. Infection and Immunity. 2013 July 11;81(8):2788–99.

17. Coleman SA, Minnick MF. Establishing a direct role for the Bartonella bacilliformis invasion-associated locus B (IalB) protein in human erythrocyte parasitism. Infect Immun. 2001 July;69(7):4373–81.

18. Dorison L, Béchon N, Martin-Gallausiaux C, Chamorro-Rodriguez S, Vitrenko Y, Ouazahrou R, et al. Identification of Veillonella parvula and Streptococcus gordonii adhesins mediating co-aggregation and its impact on physiology and mixed biofilm structure. mBio. 2024 Nov 11;15(12):e02171–24.

19. Witwinowski J, Sartori-Rupp A, Taib N, Pende N, Tham TN, Poppleton D, et al. An ancient divide in outer membrane tethering systems in bacteria suggests a mechanism for the diderm-to-monoderm transition. Nat Microbiol. 2022 Mar;7(3):411–22.

20. Silale A, Zhu Y, Witwinowski J, Smith RE, Newman KE, Bhamidimarri SP, et al. Dual function of OmpM as outer membrane tether and nutrient uptake channel in diderm Firmicutes. Nat Commun. 2023 Nov 6;14(1):7152.

21. Beaud Benyahia B, Taib N, Beloin C, Gribaldo S. Terrabacteria: redefining bacterial envelope diversity, biogenesis and evolution. Nat Rev Microbiol. 2025 Jan;23(1):41–56.

22. Rooke JL, Icke C, Wells TJ, Rossiter AE, Browning DF, Morris FC, et al. BamA and BamD Are Essential for the Secretion of Trimeric Autotransporter Adhesins. Frontiers in Microbiology [Internet]. 2021 [cited 2022 Oct 19];12. Available from: https://www.frontiersin.org/articles/10.3389/fmicb.2021.628879

23. Sikdar R, Bernstein HD. Sequential Translocation of Polypeptides across the Bacterial Outer Membrane through the Trimeric Autotransporter Pathway. mBio. 2019 Oct 22;10(5):e01973–19.

24. Kumar S, Inns PG, Ward S, Lagage V, Wang J, Kaminska R, et al. Immobile lipopolysaccharides and outer membrane proteins differentially segregate in growing Escherichia coli. Proceedings of the National Academy of Sciences. 2025 Mar 11;122(10):e2414725122.

25. Cao P, Wall D. The Fluidity of the Bacterial Outer Membrane Is Species Specific. BioEssays. 2020;42(8):1900246.

26. Mamou G, Corona F, Cohen-Khait R, Housden NG, Yeung V, Sun D, et al. Peptidoglycan maturation controls outer membrane protein assembly. Nature. 2022 June;606(7916):953–9.

27. Knapp S, Brodal C, Peterson J, Qi F, Kreth J, Merritt J. Natural Competence Is Common among Clinical Isolates of Veillonella parvula and Is Useful for Genetic Manipulation of This Key Member of the Oral Microbiome. Front Cell Infect Microbiol [Internet]. 2017 [cited 2020 Dec 16];7. Available from: https://www.frontiersin.org/articles/10.3389/fcimb.2017.00139/full?report=reader

28. Danne C, Guérillot R, Glaser P, Trieu-Cuot P, Dramsi S. Construction of isogenic mutants in Streptococcus gallolyticus based on the development of new mobilizable vectors. Research in Microbiology. 2013 Dec;164(10):973–8.

29. Zhou P, Liu J, Merritt J, Qi F. A YadA-like autotransporter, Hag 1, in Veillonella atypica is a Multivalent Hemagglutinin Involved in Adherence to Oral Streptococci, Porphyromonas gingivalis, and Human Oral Buccal Cells. Mol Oral Microbiol. 2015 Aug;30(4):269–79.

30. Shen W, Ren H. TaxonKit: A practical and efficient NCBI taxonomy toolkit. Journal of Genetics and Genomics. 2021 Sept 20;48(9):844–50.

31. Ludwiczak J, Winski A, Szczepaniak K, Alva V, Dunin-Horkawicz S. DeepCoil-a fast and accurate prediction of coiled-coil domains in protein sequences. Bioinformatics. 2019 Aug 15;35(16):2790–5.

32. Evans R, O’Neill M, Pritzel A, Antropova N, Senior A, Green T, et al. Protein complex prediction with AlphaFold-Multimer [Internet]. bioRxiv; 2022 [cited 2024 June 5]. p. 2021.10.04.463034. Available from: https://www.biorxiv.org/content/10.1101/2021.10.04.463034v2

33. Pettersen EF, Goddard TD, Huang CC, Meng EC, Couch GS, Croll TI, et al. UCSF ChimeraX: Structure visualization for researchers, educators, and developers. Protein Sci. 2021 Jan;30(1):70–82.

34. Miele V, Penel S, Duret L. Ultra-fast sequence clustering from similarity networks with SiLiX. BMC Bioinformatics. 2011 Apr 22;12(1):116.

35. Minh BQ, Schmidt HA, Chernomor O, Schrempf D, Woodhams MD, von Haeseler A, et al. IQ-TREE 2: New Models and Efficient Methods for Phylogenetic Inference in the Genomic Era. Molecular Biology and Evolution. 2020 May 1;37(5):1530–4.

36. Letunic I, Bork P. Interactive Tree Of Life (iTOL) v4: recent updates and new developments. Nucleic Acids Research. 2019 July 2;47(W1):W256–9.

37. Coleman GA, Davín AA, Mahendrarajah TA, Szánthó LL, Spang A, Hugenholtz P, et al. A rooted phylogeny resolves early bacterial evolution. Science. 2021 May 7;372(6542):eabe0511.

38. Schindelin J, Arganda-Carreras I, Frise E, Kaynig V, Longair M, Pietzsch T, et al. Fiji: an open-source platform for biological-image analysis. Nat Methods. 2012 July;9(7):676–82.

