## Supplementary Material for "Global analysis of trimeric autotransporters reveals phylogenetically restricted secretion mechanism adaptations"

### Supplementary TABLE S1: Strains used in this study

| Name | Genotype | Source |
|---|---|---|
| ***V. parvula SKV38*** | | |
| WT | - | (Knapp et al. 2017) |
| Δ*vtaA* | Δ*vtaA ::*tetM | (Béchon et al. 2020) |
| *vtaA*ΔCC | *vtaA*Δ8935:9123 ::tetM | This study |
| *vtaA*ΔCC-HA | *vtaA*Δ8935:9123-HA ::-tetM | This study |
| *vtaA*-HA | *vtaA*-HA ::-tetM | This study |
| *vtaE*ΔCC-HA | *vtaE*Δ9253:9426-HA::tetM | This study |
| *vtaF*ΔCC-HA | *vtaF*Δ9445:9576-HA::tetM | This study |
| *vtaE*ΔCC | *vtaE*Δ9253:9426::tetM | This study |
| *vtaF*ΔCC | *vtaF*Δcoiled-coi:tetM | This study |
| *vtaA-CC*$_{vtaF}$ | *vtaA*(1:8865)-*vtaF*(9381:9576)::tetM | This study |
| *vtaA-SLH*$_{00037}$ | *vtaA*(1:8865)-FNLLGLLA_00037(528:969)::tetM | This study |
| *vtaA-SLH*$_{vtaH}$ | *vtaA*(1:8865)-*vtaH*(4512:4959)::tetM | This study |
| Δ*bfvRS* | Δ*bfvRS*::catp | Audrain et al. In prep |
| Δ*bfvRS VtaG-HA*$_{Cter}$ | Δ*bfvRS*::catp, VtaG-HA::tetM | This study |
| Δ*bfvRS VtaG*Δ*SLH* | Δ*bfvRS*::catp, VtaGΔ1240:1731::tetM | This study |
| Δ*bfvRS VtaG*Δ*SLH-HA* | Δ*bfvRS*::catp, VtaGΔ1240:1731-HA::tetM | This study |
| *vtaA-25pCC* | VtaA-Δ8985:9123-HA::tetM | This study |
| *vtaA-50pCC* | VtaA-Δ9030:9123-HA::tetM | This study |
| *vtaA-75pCC* | VtaA-Δ9075:9123-HA::tetM | This study |
| Δ*bfvRS vtaG*-TRYE-HA | Δ*bfvRS*::catp, VtaG-TRYE$_{496}$AAAA-HA::tetM | This study |
| Δ*bfvRS vtaG*-WAY-HA | Δ*bfvRS*::catp, VtaG-WAY$_{467}$AAA-HA::tetM | This study |
| Δ*bfvRS* Δ*vtaF* Δ*vtaG* | Δ*bfvRS*:catP, ΔvtaF::kanR, ΔVtaG::eryR | This study |
| Δ*bfvRS* Δ*vtaF* | Δ*bfvRS*:catP, ΔvtaF::kanR, | This study |
| Δ*bfvRS* Δ*vtaF HA*$_{nter}$-*vtaG* | HA(nter)-VtaG::tetM, Δ*bfvRS*:catP, ΔvtaF::kanR | This study |
| Δ*bfvRS* Δ*vtaF HA*$_{nter}$-*vtaG*Δ*SLH* | HA(nter)-VtaGΔ1240:1731::tetM, Δ*bfvRS*:catP, ΔvtaF::kanR | This study |
| Δ*bfvRS* Δ*vtaF HA*$_{nter}$-*vtaG*-TRYE | HA(nter)-VtaG-TRYE$_{496}$AAAA::tetM, Δ*bfvRS*:catP, ΔvtaF::kanR | This study |
| Δ*bfvRS* Δ*vtaF HA*$_{nter}$-*vtaG-CCvtaA* | HA(nter)-VtaG-VtaA_cter::tetM, Δ*bfvRS*:catP, ΔvtaF::kanR | This study |
| Δ*bfvRS* Δ*vtaF vtaG-CCvtaA* | VtaG-VtaA_cter_V2::tetM, Δ*bfvRS*:catP, ΔvtaF::kanR | This study |
| Δ*bfvRS* Δ*vtaF HA*$_{nter}$-*vtaG-WAY* | HA(nter)-VtaG-WAY$_{467}$AAA::tetM, Δ*bfvRS*::catP, ΔvtaF::kanR | This study |
| Δ*bfvRS* Δ*vtaF HA*$_{nter}$-*vtaG-WAY-TRYE* | HA(nter)-VtaG-WAY$_{467}$AAA-TRYE$_{496}$AAAA::tetM, Δ*bfvRS*::catP, ΔvtaF::kanR | This study |
| *Escherichia coli* DH5α | pET22b-VtaA | This study |

| | | |
|---|---|---|
| *Escherichia coli BL21de3* | pDIA17 - pET22b-vtaA | This study |
| *Streptococcus oralis* ATCC10557 | WT | |
| *Streptococcus gordonii* DL1 | WT | |
| Plasmids | pET22b-vtaA(133:2100) | This study |

### Supplementary Table S2: primers used in this study

| Name | sequence ( 5' -> 3' ) | construction |
|---|---|---|
| 5F_VtaA_HA | gctgcaattgatagaaatgctgctg | vtaA-HA |
| 5R_vtaA_tetR_HA | cttcactcgcctgcattttactctcttaacctttagaagcta | vtaA-HA |
| TetMF | AGTAAAATGCAGGCGAGTGAAG | cassette tetracycline resistance |
| TetM R | GTGGATCCACAGGACACAAT | cassette tetracycline resistance |
| 3F_VtaA_TetR | attgtgtcctgtggatccacttgagaactaattttgata | 3' homology of *vtaA* for tetM |
| 3R_VtaA_TetR | ggcctttcaaaagtccatttcg | 3' homology of *vtaA* for tetM |
| 5R_VtaA_coiledcoiled_tetR | cttcactcgcctgcattttactctcTCATTAtacataaacagagctaataggccc | vtaA-ΔCC |
| F_VtaA_His | CTTTAAGAAGGAGATATACATATGaatgctagcatctctatggcgg | *cloning of vtaA in pet22B* |
| R_Vta_His | AGTGGTGGTGGTGGTGGTGagtatttttatctattgcagttttattgtcag | *cloning of vtaA in pet22B* |
| pET22b_LINEAR_F | CACCACCACCACCACCACT | *cloning of vtaA in pet22B* |
| pET22Bb_LINEAR_R | ATGTATATCTCCTTCTTAAAGTTAAACAAAATTATTTC | *cloning of vtaA in pet22B* |
| 5R_VtaA_coiledcoiled_HA | ataatctggaacatcatatggatagctgccgctaccgctacctacataaacagagctaataggccccgc | vtaA-ΔCC-HA |
| HACter_TetR_F | catatgatgttccagattatgctTAAgagagtaaaatgcaggcgagtgaag | *tetM with a HA-tag* |
| VtaA_5R_VtaA_HACter | ataatctggaacatcatatggatagctgccgctaccgctaccacctttagaagctaataattgatttactacgg | *tetM with a HA-tag and a linker* |
| 5F_VtaF_coiled-coil_tetR | gactaagagaacaagcacaactg | vtaF-ΔCC |
| 5R_VtaF_coiledcoil-tetR | cttcactcgcctgcattttactctcttatgtagaactgataggtcctttac | vtaF-ΔCC |
| 3F_VtaF_tetR | attgtgtcctgtggatccactacgttcatactgaacattacgg | vtaF-ΔCC |
| 3R_VtaF_tetR | tttgcacaatttcacggcc | vtaF-ΔCC |
| 5R-VtaF-coilcoiled-HA | agcataatctggaacatcatatggatagctgccgctaccgctacctgtagaactgataggtcctttac | vtaF-ΔCC-HA |
| 5F_VtaE_coiled-coil_tetR | gaaaggtatggattttgccg | vtaE-ΔCC |
| 5R_VtaE_coiled-coil_tetR | cttcactcgcctgcattttactctcttagtacatagagctgataggac | vtaE-ΔCC |
| 3F_VtaE_tetR | attgtgtcctgtggatccacttcacatctgtttcatcttatattc | vtaE-ΔCC |
| 3R_VtaE_tetR | caatagcaccgatttgttc | vtaE-ΔCC |
| 5R_VtaE_HA_coilcoiled | agcataatctggaacatcatatggatagctgccgctaccgctaccgtacatagagctgataggac | vtaE-ΔCC-HA |
| R_vtaF_domain_swap | tcactcgcctgcattttactctcTTATAAACCTAGTCTTGCCATCATTG | vtaA-CCvtaF |
| R_vtaH_domain_swap | tcactcgcctgcattttactctcTTAACCGCGACCTTTAATAAC | vtaA-SLHvtaH |
| R_00037_domain_swap | tcactcgcctgcattttactctcCTATTTTTTATTGACCCTTACACGT | vtaA-SLH00037 |

| | | |
|---|---|---|
| 5R_vtaA_dom_swap | accaaatttatgcgttacacc | vtaA-cter |
| F_vtaH_dom_swap | gtgtaacgcataaatttggtCAAGGCAATCATGTGTC | vtaA-SLHvtaH |
| F_vtaF_dom_swap | gtgtaacgcataaatttggtAACAGAGACGGCGAAG | vtaA-CCvtaF |
| F_00037_dom_swap | gtgtaacgcataaatttggtGCTCACAGTAATGTATCTC | vtaA-SLH00037 |
| 5R_VtaG_HA_linker | ctggaacatcatatggatagctgccgctaccgctaccttgattattgtatatacggg aacc | *vtaG-HA-cter* |
| 3F_VtaG_tetM | attgtgtcctgtggatccacaattaacttttctagagtttgtaaaaactc | *vtaG-HA-cter* |
| 5F_VtaG | gcaacacatttaaactctatcgc | *vtaG-HA-cter* |
| 5R_VtaG_dSLH-HA | taatctggaacatcatatggatagctgccgctaccgctaccaggggaacgtttac ctagg | *vtaGΔSLH-HA-cter* |
| 5R_VtaG_dSLH-tetM | cttcactcgcctgcattttactctccTAaggggaacgtttacctag | *vtaGΔSLH* |
| 3R_RasP | ccgtaatgtatcaccaatgc | *ΔrasP* |
| 5F_RasP | ggctggtcttgtagatgaag | *ΔrasP* |
| 3F_Δ_RasP_kana | TTTACTGGATGAATTGTTTTAGgacagatgggagattacaatg | *ΔrasP* |
| 5R_Δ_RasP_kana | CCGGTGATATTCTCATTTTAGCCATatattctcctataacatcaac gatg | *ΔrasP* |
| 3R_ctpB | gccttccataccagatacatc | *ΔctpB* |
| 5F_ctpB | cggtcgtttaagttacttagatg | *ΔctpB* |
| 5R_Δ_ctpB_kana | CCGGTGATATTCTCATTTTAGCCATattataaatatcccataggat ccac | *ΔctpB* |
| 3F_Δ_ctpB_kana | TACTGGATGAATTGTTTTAGtgcataggagttaaattatgtttatagat ag | *ΔctpB* |
| 3R_bepA | gtatactccatcatagcctcc | *ΔbepA* |
| 5F_bepA | ccatggatattggtatacttaggc | *ΔbepA* |
| 3F_Δ_bepA_kana | ATTTTACTGGATGAATTGTTTTAGgagctatacatggcatataattt gc | *ΔbepA* |
| 5R_ΔbepA_kana | cttcactcgcctgcattttatctctcaacctctcataatgataaatatac | *ΔbepA* |
| 3F_Δ_hhoB_kana | ATTTTACTGGATGAATTGTTTTAGaatctagtaaataaaacttaatt aagtattcc | *ΔhhoB* |
| 5F_hhoB | caatataaacgtttaacagagc | *ΔhhoB* |
| 5R_Δ_hhob_kana | cttcactcgcctgcattttagtagatctccttatattcttcttaatattc | *ΔhhoB* |
| 3R_hhoB | ccccggtaattgaggtaaaagac | *ΔhhoB* |
| R_VtaE_HA | atagctgccgctaccgctacccatgcctaaacgactcataataag | *vtaE*-HA |
| F_linker_HA | gtagcggtagcggcagctat | different Cter-VtaA |
| 5R_vtaA_25CC_HA | ggatagctgccgctaccgctacccgtttttgattactattttctttttg | *vtaA*-25pCC |
| 5R_vtaA_50CC_HA | ggatagctgccgctaccgctaccttcagcttctaatgtatttagacgc | *vtaA*-50pCC |
| 5R_vtaA_75CC_HA | ggatagctgccgctaccgctacctaagccttgtttagtttcagcc | *vtaA*-75pCC |
| F_VtaG_W467A | gcaaccacGCAgcAGCTcaatatattagccaattgg | vtaG-WAY |
| R_VtaG_W467A | atattgAGCTgcTGCgtggttgcgaggaacgtc | vtaG-WAY |
| F_VtaG_TRYE496A AAA | gatgGCAGCTGCAGCTttcgctaccatgttgtaccg | vtaG-TRYE |
| R_VtaG_TRYE496A AAA | gcgaaAGCTGCAGCTGCcatcttaacatcgcctttga | vtaG-TRYE |
| 5R_VtaG_HA_nter | ataatctggaacatcatatggataagcagcgtctactggattaa | vtaG-HAnter |
| 3F_VtaG_HA_nter | ccatatgatgttccagattatgcttctgttgtaatggacaacac | vtaG-HAnter |
| 3R_VtaG-tetM | cttcactcgcctgcattttactctcctattgattattgtatatacgggaacc | vtaG-HAnter |
| F_VtaA_for_vtaG | aggtgtatctattcgcctaggtggtagtcgtgaaagaaaagacg | vtaG-CCvtaA |
| R_VtaG_for_VtaA_C C | acctaggcgaatagatacac | vtaG-CCvtaA |

### Supplementary Table S3: most abundant functions identified by HMMscan in the adhesins and their neighbors.

| Pfam name | function | occurences | Pfam |
|---|---|---|---|
| YadA_head | YadA head domain repeat (2 copies) | 35072 | PF05658.17 |
| YadA_stalk | Coiled stalk of trimeric autotransporter adhesin | 18117 | PF05662.17 |
| YadA_anchor | YadA-like membrane anchor domain | 3171 | PF03895.18 |
| ESPR | Extended Signal Peptide of Type V secretion system | 1285 | PF13018.9 |
| Response_reg | Response regulator receiver domain | 758 | PF00072.27 |
| OmpA | OmpA family | 612 | PF00691.23 |
| HATPase_c | Histidine kinase-, DNA gyrase B-, and HSP90-like ATPase | 439 | PF02518.29 |
| SmpA_OmlA | SmpA / OmlA family | 437 | PF04355.16 |
| MFS_1 | Major Facilitator Superfamily | 423 | PF07690.19 |
| HTH_18 | Helix-turn-helix domain | 416 | PF12833.10 |
| ABC_tran | ABC transporter | 372 | PF00005.30 |
| HTH_AraC | Bacterial regulatory helix-turn-helix proteins, AraC family | 349 | PF00165.26 |
| GerE | Bacterial regulatory proteins, luxR family | 332 | PF00196.22 |
| DUF2827 | Protein of unknown function (DUF2827) | 314 | PF10933.11 |
| Trans_reg_C | Transcriptional regulatory protein, C terminal | 283 | PF00486.31 |
| LysR_substrate | LysR substrate binding domain | 281 | PF03466.23 |
| HTH_1 | Bacterial regulatory helix-turn-helix protein, lysR family | 277 | PF00126.30 |
| HisKA | His Kinase A (phospho-acceptor) domain | 258 | PF00512.28 |
| Sugar_tr | Sugar (and other) transporter | 238 | PF00083.27 |
| Peptidase_S8 | Subtilase family | 194 | PF00082.25 |
| IalB | Invasion associated locus B (IalB) protein | 184 | PF06776.15 |
| TPR_16 | Tetratricopeptide repeat | 184 | PF13432.9 |
| Trp_ring | Trimeric autotransporter adhesin Trp ring domain | 174 | PF18669.4 |
| BPD_transp_1 | Binding-protein-dependent transport system inner membrane component | 170 | PF00528.25 |
| Acetyltransf_1 | Acetyltransferase (GNAT) family | 162 | PF00583.28 |
| TPR_8 | Tetratricopeptide repeat | 151 | PF13181.9 |
| Aldedh | Aldehyde dehydrogenase family | 150 | PF00171.25 |
| Collagen | Collagen triple helix repeat (20 copies) | 148 | PF01391.21 |
| TonB_dep_Rec | TonB dependent receptor | 145 | PF00593.27 |
| Plug | TonB-dependent Receptor Plug Domain | 145 | PF07715.18 |
| EamA | EamA-like transporter family | 143 | PF00892.23 |
| DUF4189 | Domain of unknown function (DUF4189) | 138 | PF13827.9 |
| T2SSF | Type II secretion system (T2SS), protein F | 137 | PF00482.26 |
| AAA | ATPase family associated with various cellular activities (AAA) | 137 | PF00004.32 |
| PPC | Bacterial pre-peptidase C-terminal domain | 135 | PF00106.28 |
| adh_short | short chain dehydrogenase | 135 | PF04151.18 |
| EAL | EAL domain | 134 | PF00563.23 |
| adh_short_C2 | Enoyl-(Acyl carrier protein) reductase | 133 | PF13561.9 |
| Sel1 | Sel1 repeat | 129 | PF08238.15 |
| Acetyltransf_7 | Acetyltransferase (GNAT) domain | 121 | PF13508.10 |
| Methyltransf_25 | Methyltransferase domain | 118 | PF13649.9 |
| Helicase_C | Helicase conserved C-terminal domain | 115 | PF00271.34 |
| SLH | S-layer homology domain | 114 | PF00395.23 |
| HTH_3 | Helix-turn-helix | 113 | PF01381.25 |
| OEP | Outer membrane efflux protein | 112 | PF02321.21 |
| Methyltransf_31 | Methyltransferase domain | 110 | PF08659.13 |
| KR | KR domain | 110 | PF13847.9 |
| Methyltransf_11 | Methyltransferase domain | 109 | PF08241.15 |
| PAS_4 | PAS fold | 108 | PF08448.13 |
| Abhydrolase_1 | alpha/beta hydrolase fold | 107 | PF00561.23 |
| Abhydrolase_6 | Alpha/beta hydrolase family | 106 | PF12697.10 |
| AIRS_C | AIR synthase related protein, C-terminal domain | 105 | PF02769.25 |
| AAA_5 | AAA domain (dynein-related subfamily) | 103 | PF07728.17 |

| | | | |
|---|---|---|---|
| Sigma70_r4_2 | Sigma-70, region 4 | 102 | PF08281.15 |
| SBP_bac_3 | Bacterial extracellular solute-binding proteins, family 3 | 100 | PF00497.23 |
| TPR_10 | Tetratricopeptide repeat | 94 | PF13374.9 |
| tRNA-synt_1 | tRNA synthetases class I (I, L, M and V) | 94 | PF00133.25 |
| T2SSE | Type II/IV secretion system protein | 90 | PF00437.23 |
| Phage_integrase | Phage integrase family | 90 | PF00589.25 |
| Sigma70_r4 | Sigma-70, region 4 | 89 | PF04545.19 |

### Supplementary Table S4: most abundant functions identified by HMMscan in the adhesins

| target | target_description | count |
|---|---|---|
| YadA_head | YadA head domain repeat (2 copies) | 32242 |
| YadA_stalk | Coiled stalk of trimeric autotransporter adhesin | 16463 |
| YadA_anchor | YadA-like membrane anchor domain | 3011 |
| ESPR | Extended Signal Peptide of Type V secretion system | 1091 |
| Trp_ring | Trimeric autotransporter adhesin Trp ring domain | 161 |
| Collagen | Collagen triple helix repeat (20 copies) | 136 |
| SLH | S-layer homology domain | 71 |
| TAA-Trp-ring | Tryptophan-ring motif of head of Trimeric autotransporter adhesin | 53 |
| CFSR | Collagen-flanked surface repeat | 39 |
| OmpA | OmpA family | 13 |
| HiaBD2 | HiaBD2_N domain of Trimeric autotransporter adhesin (GIN) | 8 |
| UspA1_rep | Ubiquitous surface protein adhesin repeat | 6 |
| Adeno_shaft | Adenoviral fibre protein (repeat/shaft region) | 5 |
| DUF285 | Mycoplasma protein of unknown function, DUF285 | 3 |
| DUF1079 | Repeat of unknown function (DUF1079) | 3 |
| Alanine_zipper | Alanine-zipper, major outer membrane lipoprotein | 2 |
| Av_adeno_fibre | Avian adenovirus fibre, N-terminal | 1 |

### Supplementary Table S5: List of all TAAs in *V. parvula*

| name | gene | size (a.a) | periplasmic domain | | function |
|---|---|---|---|---|---|
| | | | type | size (a.a) | |
| VtaA | FNLLGLLA_00516 | 3041 | coiled-coil | 86 | autoaggregation / coaggregation *S. oralis* |
| VtaB | FNLLGLLA_00034 | 470 | coiled-coil | 45 | - |
| VtaC | FNLLGLLA_00038 | 2811 | coiled-coil | 74 | - |
| VtaD | FNLLGLLA_00044 | 2072 | coiled-coil | 59 | coaggregation *S. gordonii* |
| VtaE | FNLLGLLA_00045 | 3142 | coiled-coil | 79 | coaggregation *S. gordonii* |
| VtaF | FNLLGLLA_00046 | 3193 | coiled-coil | 67 | Biofilm formation / Surface attachment |
| VtaG | FNLLGLLA_00098 | 577 | SLH (455-518) | 170 | Biofilm formation |
| VtaH | FNLLGLLA_00099 | 1653 | SLH (1550-1613) | 151 | - |
| VtaI | FNLLGLLA_0001790 | 833 | SLH (705-768) | 175 | - |

**Supplementary text 1: does the SLH domain of TAAs bind PG?**

TAAs in Terrabacteria often possess S-layer homology (SLH) domain that were described to bind to PG, sometimes modified with aliphatic polyamines such as cadaverine or putrescine. The other main protein with an SLH domain is OmpM, the main porin and membrane tethering system, which trimerize, similarly to TAAs, with each monomer bringing one SLH domain (Witwinowski et al. 2022; Silale et al. 2023). Additionally, SLH domains in TAAs possess the conserved residues important for SLH binding and we therefore think that these SLH domains are indeed binding peptidoglycan. We first tried to prove this experimentally by purifying the SLH domain of VtaH, incubating it with different concentrations of purified *Veillonella* PG and retrieving the supernatant of the centrifuged mixture. We could not see a gradual reduction of the purified protein quantity as we would have expected for PG binding. (Figure TS1). However, this strategy also had been not successful with the SLH domain of the OmpM protein, and the only way to show PG binding was to natively purify the full OmpM protein in *Veillonella* (Silale et al. 2023). VtaG is less abundant than OmpM and we haven't attempted to purify the TAA natively but instead have tried to observe PG binding binding of VtaG from crude lysat of *Veillonella*, which also did not work (Figure TS2). The only indirect proof that we have of PG binding is the fact that VtaG mutated in supposedly essential residues in the SLH domain presents a striking different localization at the cell surface.

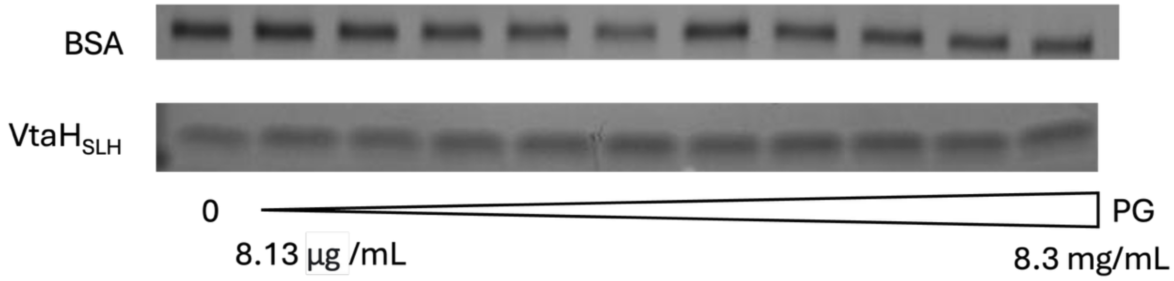

**Figure TS1: Purified SLH domain does not bind PG.** Coomassie stained SDS-page gels presenting the results of the PG binding assay of VtaH purified SLH domain. 5 µg of the purified protein or BSA was incubated with increasing (in twofold) concentrations of PG.

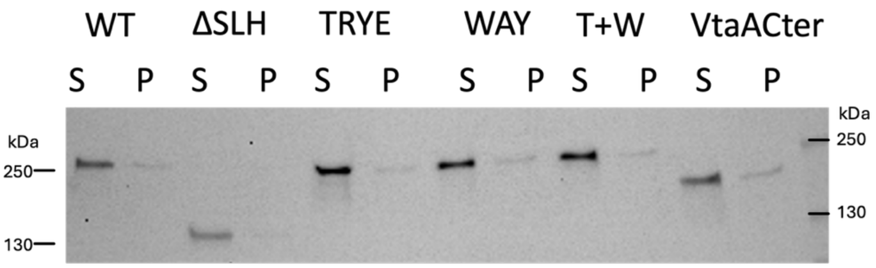

**Figure TS2**: **VtaG from a whole-cell lysat does not bind PG.** Western-blot using an anti-HA antibody recognizing an $HA_{Nter}$-tagged *vtaG* with various modifications of its SLH domain in a Δ *bfvR-S::catP*Δ*vtaF::kanR* backgroung. For each sample, a a crude cell lysate was produced, incubated with PG, centrifuged and the resulting supernatant (S) containing unbound proteins or pellet (P) was collected.

**Purification of the SLH domains**

The portion of VtaH (amino acids 1540 to 1652) coding for its SLH domain was amplified by PCR from *V. parvula* and the pET22b-HIS vector was linearized by PCR. The PCR products were then purified and annealed by Gibson reaction. Plasmids were dialyzed and transformed in electrocompetent *E. coli* DH5$\alpha$. After verification of the construct by sequencing, plasmid was purified and transformed in *E. coli* BL21(DE3)-pDIA17. After growth to $OD_{600}$ 0.4, cells were induced with 0.1 mM IPTG and grown for 3h at 37°C before harvesting. Cell pellet was frozen O/N then resuspended in Buffer A (30 mM Tris-HCl pH 7.9, 300 mM NaCl, 30 mM Imidazole) and lysed by sonication. Debris were pelleted by ultracentrifugation (50 000 g, 30 min) and supernatant run through a HisTrap 5 mL column on an AKTA Explorer (GE) against a gradient of imidazole (30-300 mM). The purified protein was assessed for purity by SDS-Page followed by SafeStain SimplyBlue™ (ThermoFisher) staining and dialyzed twice against 30 mM Tris-HCl pH 7.9, 300 mM NaCl using a SnakeSkin™ 3500 Da (ThermoFisher).

**Peptidoglycan isolation**

500 mL of *V. parvula* WT were grown overnight, centrifuged and resuspended in 5 mL of ice-cold MiliQ water, before addition to 10 mL of boiling 8% SDS. Volume was adjusted to obtain a 4% SDS concentration and boiled for 1h. The resulting solution was centrifuged (30 min, 150,000 g) at room temperature and washed in MiliQ water 6 times. The pellet was then resuspended in 4 mL of lysis buffer (trypsin 100 mg/mL, 50 mM Tris pH 7.0, 10 mM $CaCl_2$) and incubated overnight at 37°C. Sample was boiled for 15 min to inactivate the trypsine and washed three times with MiliQ water (30 min, 150 000 g). The resulting PG sacculi was lyophilized by vacuum centrifugation before resuspension in 500 µL of water by sonication.

**Peptidoglycan binding assay with the purified protein.**

Purified *V. parvula* PG was diluted 2-fold 9-12 times in water. 6.25 µL of purified protein or BSA (0.25 mg/mL) were mixed with 1.5 µL of 20X buffer (Tris 0.4 M, pH 8.0) and 2.25 µL $H_2O$ and added to 20 µL of each of the PG dilutions. Solutions were mixed by vortexing and left to incubate on the bench for 45 min before centrifugation (10 min, 13 000 g). 15 µL of the supernatant was resuspended in Laemmli buffer and run on an SDS-Page followed by SafeStain SimplyBlue™ (ThermoFisher) staining.

**Peptidoglycan binding assay from crude extract.**

*V. parvula* was grown in 50 mL SK O/N, pelleted and washed twice with 500 µL binding buffer (10 mM Tris-HCl pH 7.5, 150 mM NaCl), resuspended cells were then lysed on a FastPrep (2X 20s, 4m/s, 10 min on ice) and debris pelleted (15 min, 13 000 g, 4°C). 200 µL of the supernatant were mixed with purified PG at a final concentration of 1 mg/mL and incubated for 2 hours at room temperature. Samples were centrifuged, (15 min, 13 000 g, 4°C), a sample of the supernatant was retrieved for western blotting and the pellet was washed once with binding buffer, and the pellet was resuspended in 200 µL binding buffer and an equivalent volume retrieved for western blotting.

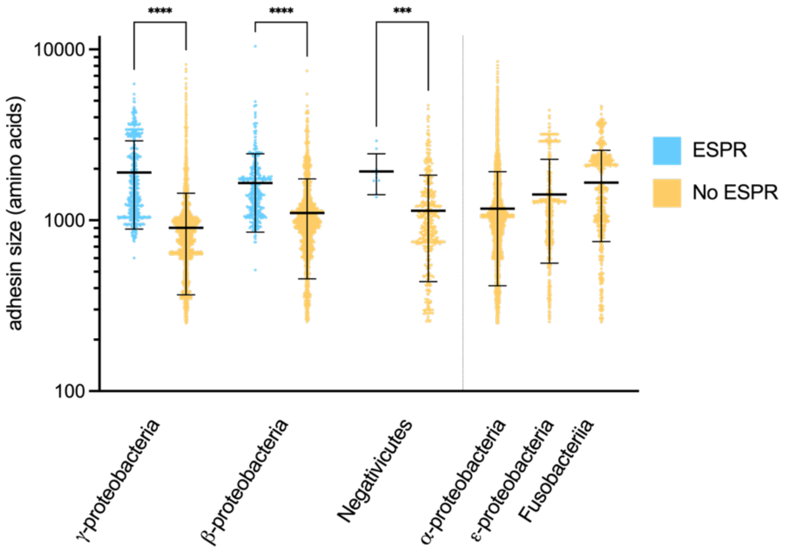

**supplementary Figure S1: Classical Va autotransporters with ESPR are on average longer than adhesins without ESPR.** Scatter plot of classical autotransporters, where each point corresponds to the size in amino acid of the adhesin. Mean and standard deviation are also represented. Adhesins were separated by class and presence or absence of ESPR. Significance of the differences between adhesins with and without ESPR was calculated using a Kruskal-Wallis test. To generate the dataset of monomeric autotransporters, the UniprotKB database was searched for proteins possessing the IPR005546 (classical autotransporter β-barrel) domain. Using customized python scripts, near duplicate proteins were dropped based on homology of their 250 last residues using the Levenstein hamming function, with a threshold of 5 residues homology, resulting in 24523 unique autotransporter sequences. Significance of the differences between adhesins with and without ESPR was calculated using a Kruskal-Wallis test (***: P ≤ 0.001, **** : P ≤ 0.0001).

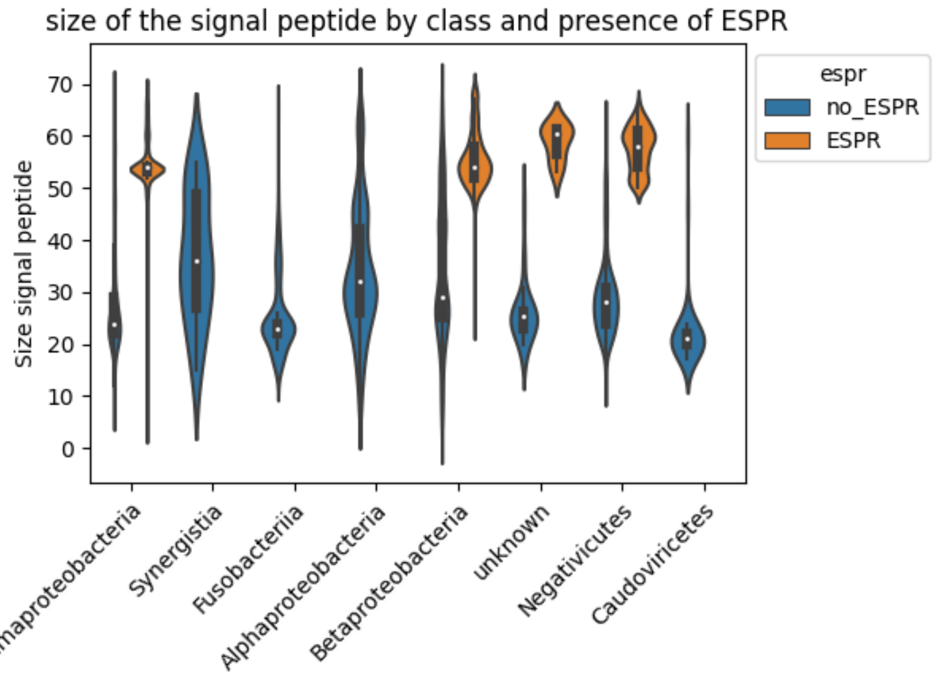

**supplementary Figure S2: Average size of signal peptides in trimeric autotransporters.** Adhesins were separated by Class and presence or absence of ESPR (PFAM13018). Presence or absence of the signal peptide was determined using SignalP 6.0 and the probable cleavage site plotted as the size of the signal peptide.

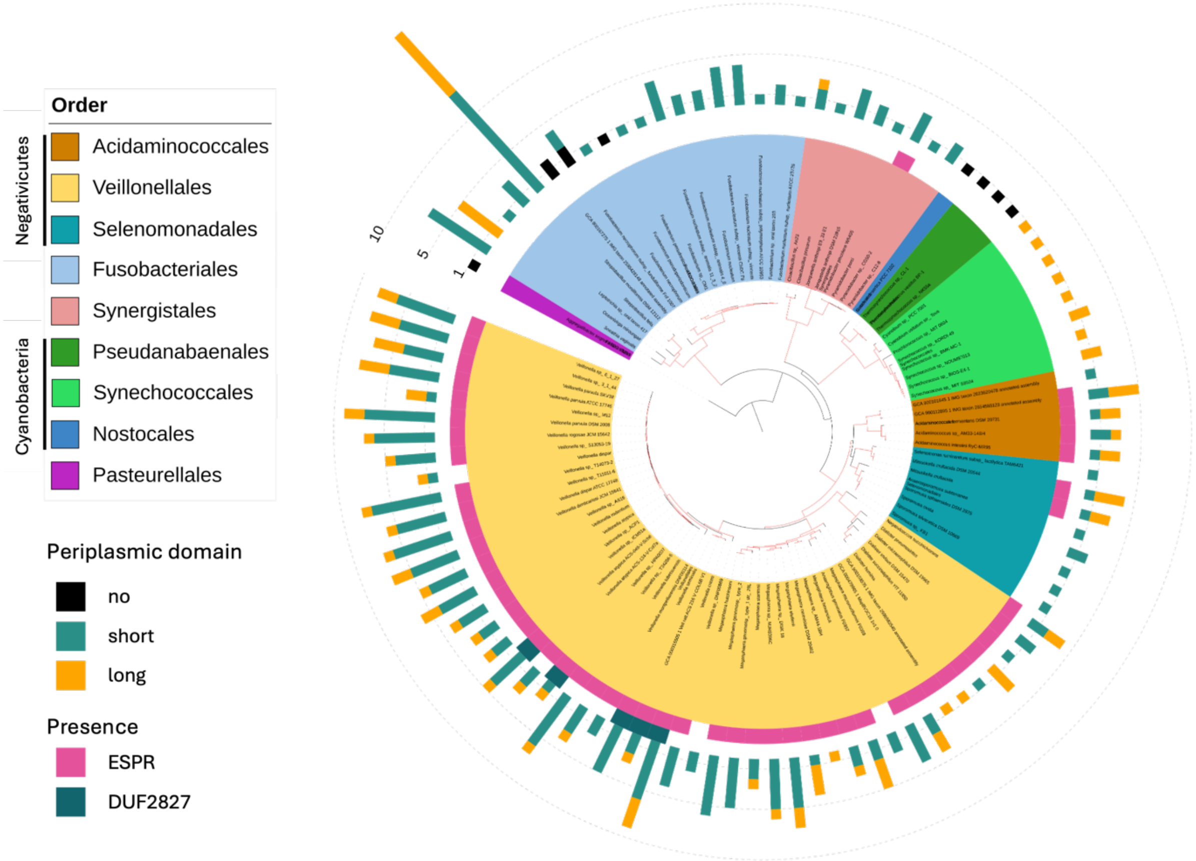

**supplementary Figure S3: Distribution of the trimeric autotransporters and their associated genes in bacteria outside of *Proteobacteria*.** We represented on a phylogeny of all selected bacteria (non Proteobacteria) the number of adhesins per genome, the type of periplasmic domain, and the presence of ESPR and DUF2827 homologues. In the inner ring, bacteria are colored by their order. On the middle rings, colored blocks represent the presence of the specified domain. The outer bars represent the number of adhesins, colored by type of periplasmic domain. Tree branches supported by bootstrap values above 0.95 are colored in red. The tree was rooted using a *Pasteurellales*, indicated in purple.

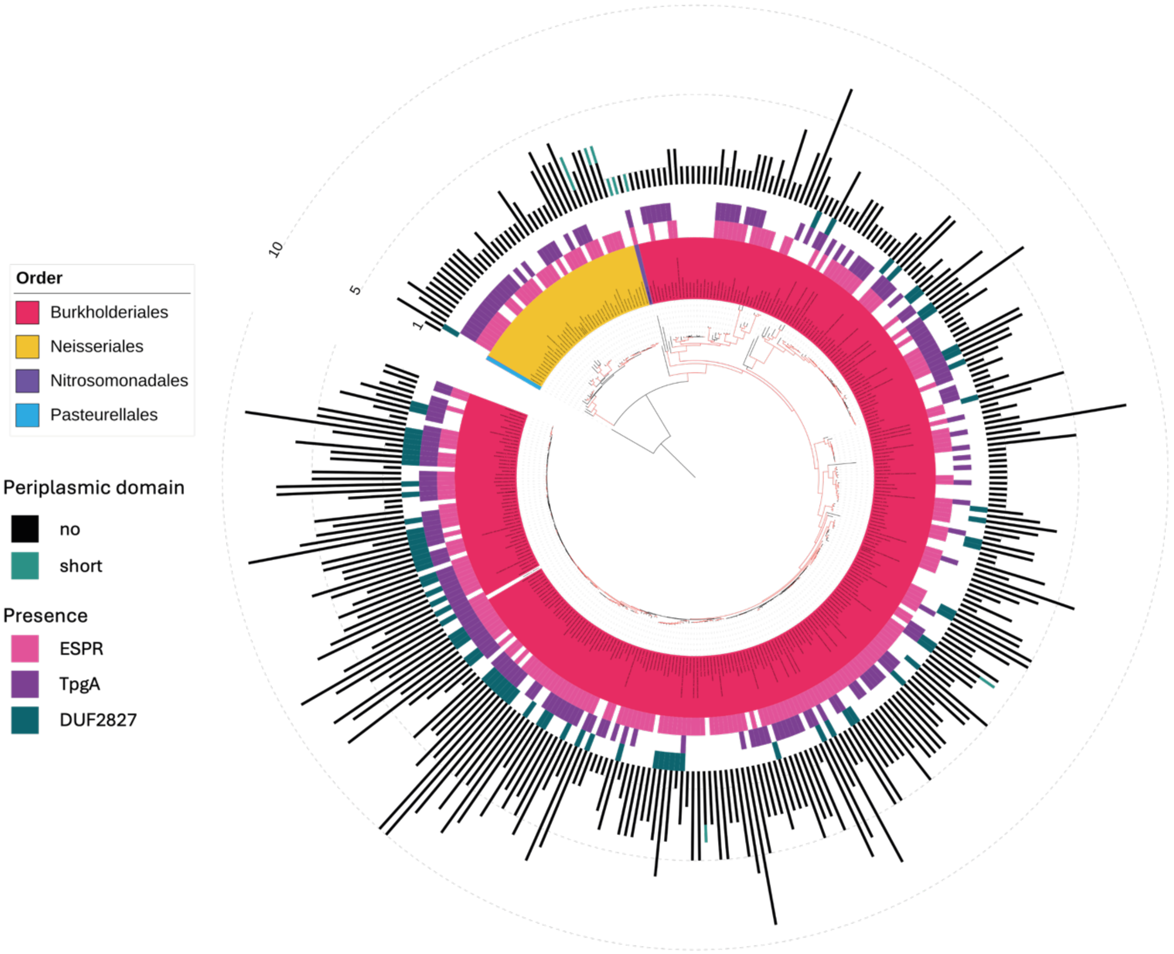

**supplementary Figure S4: Distribution of the trimeric autotransporters and their associated genes in *Betaproteobacteria*.** We represented on a phylogeny of all selected Betaproteobacteria genomes the number of adhesins per genome, the type of periplasmic domain, and the presence of ESPR, TpgA and DUF2827 homologues. In the inner ring, bacteria are colored by their order. On the middle rings, colored blocks represent the presence of the specified domain. The outer bars represent the number of adhesins, colored by type of periplasmic domain. Tree branches supported by bootstrap values above 0.95 are colored in red. The tree was rooted using a *Pasteurellales*, indicated in blue.

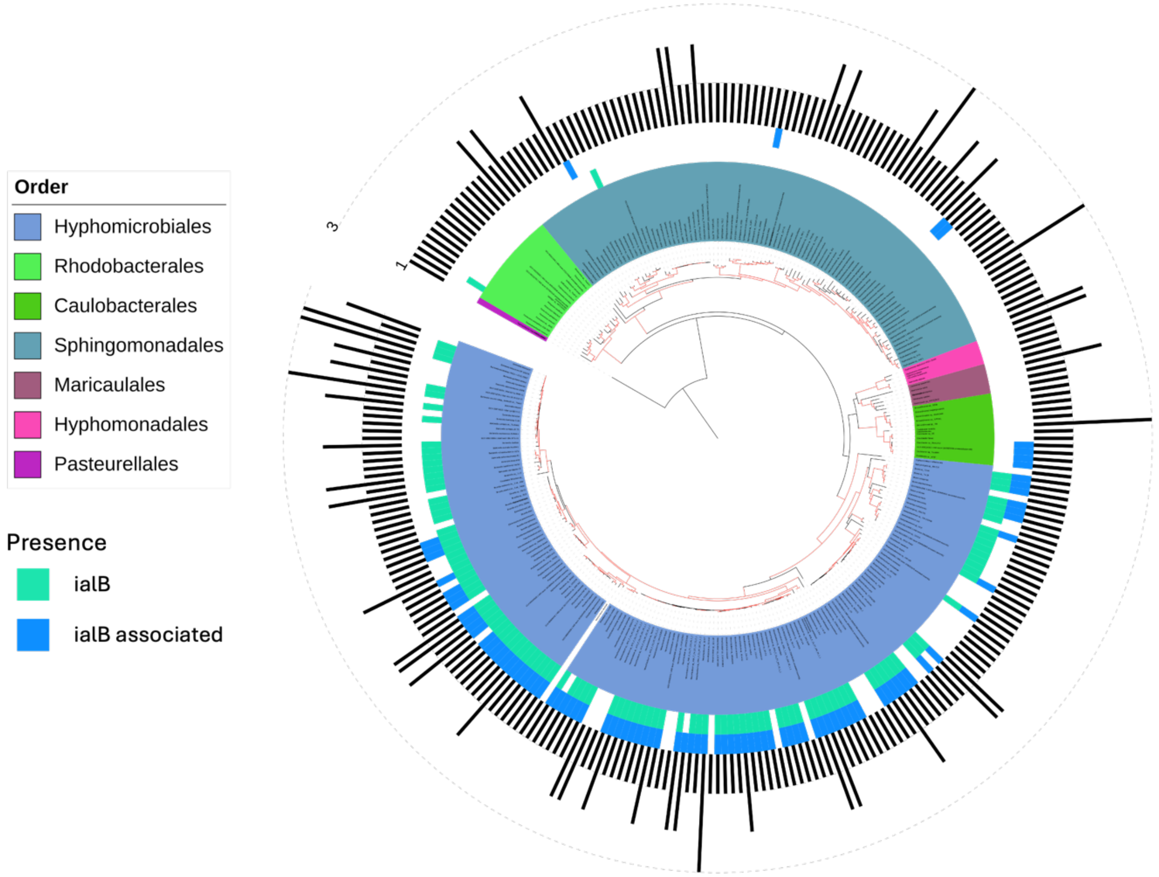

**supplementary Figure S5: Distribution of the trimeric autotransporters and their associated genes in *Alphaproteobacteria*.** We represented on a phylogeny of all selected *Alphaproteobacteria* genomes the number of adhesins per genome, the type of periplasmic domain, and the presence of IalB and IalB associated proteins homologues. In the inner ring, bacteria are colored by their order. On the middle rings, colored blocks represent the presence of the specified domain. The outer bars represent the number of adhesins, colored by type of periplasmic domain. Tree branches supported by bootstrap values above 0.95 are colored in red. The tree was rooted using a *Pasteurellales*, indicated in purple.

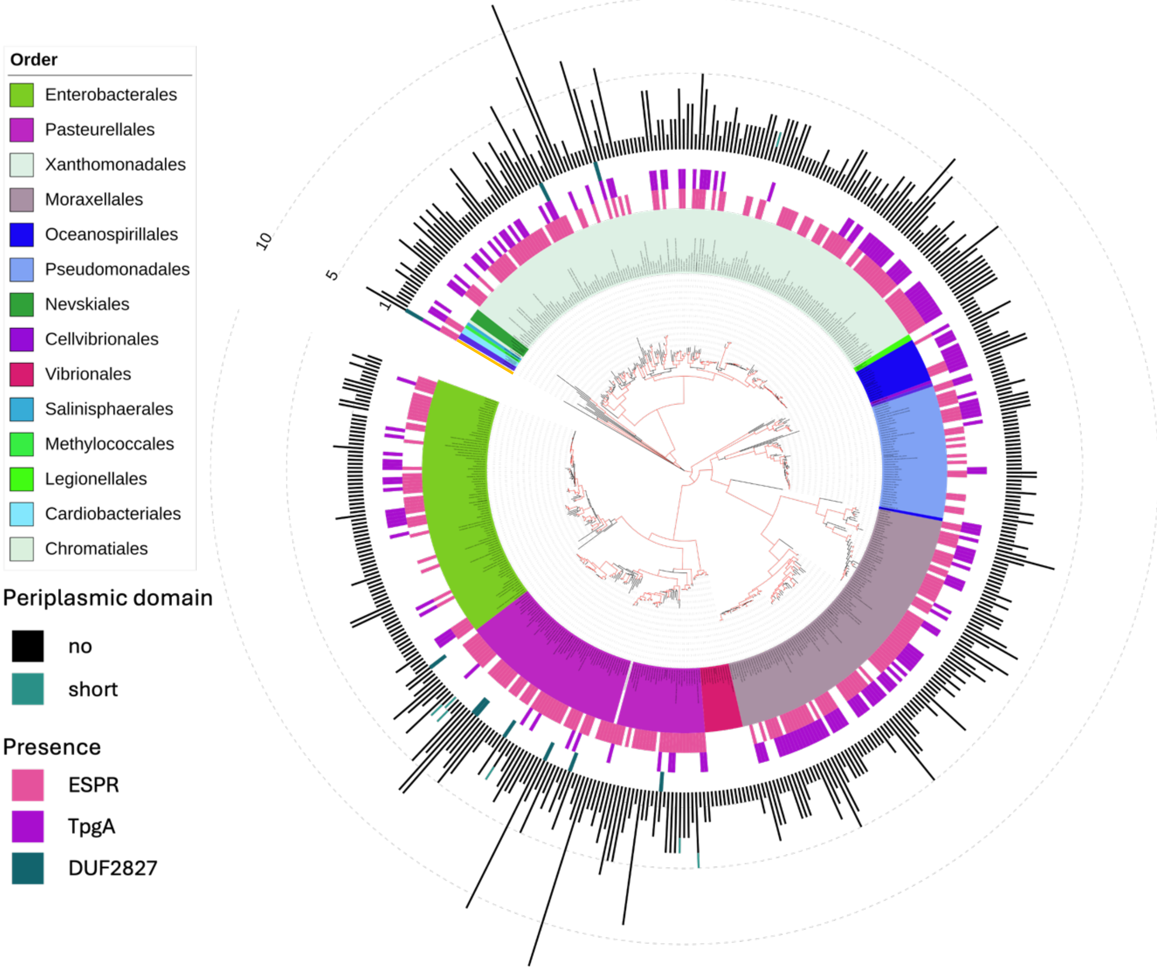

**supplementary Figure S6: Distribution of the trimeric autotransporters and their associated genes in *Gammaproteobacteria*.** We represented on a phylogeny of all selected *Gammaproteobacteria* genomes the number of adhesins per genome, the type of periplasmic domain, and the presence of ESPR, TpgA and DUF2827 homologues. In the inner ring, bacteria are colored by their order. On the middle rings, colored blocks represent the presence of the specified domain. The outer bars represent the number of adhesins, colored by type of periplasmic domain. Tree branches supported by bootstrap values above 0.95 are colored in red. The tree was rooted using a *burkholderia*, indicated in orange.

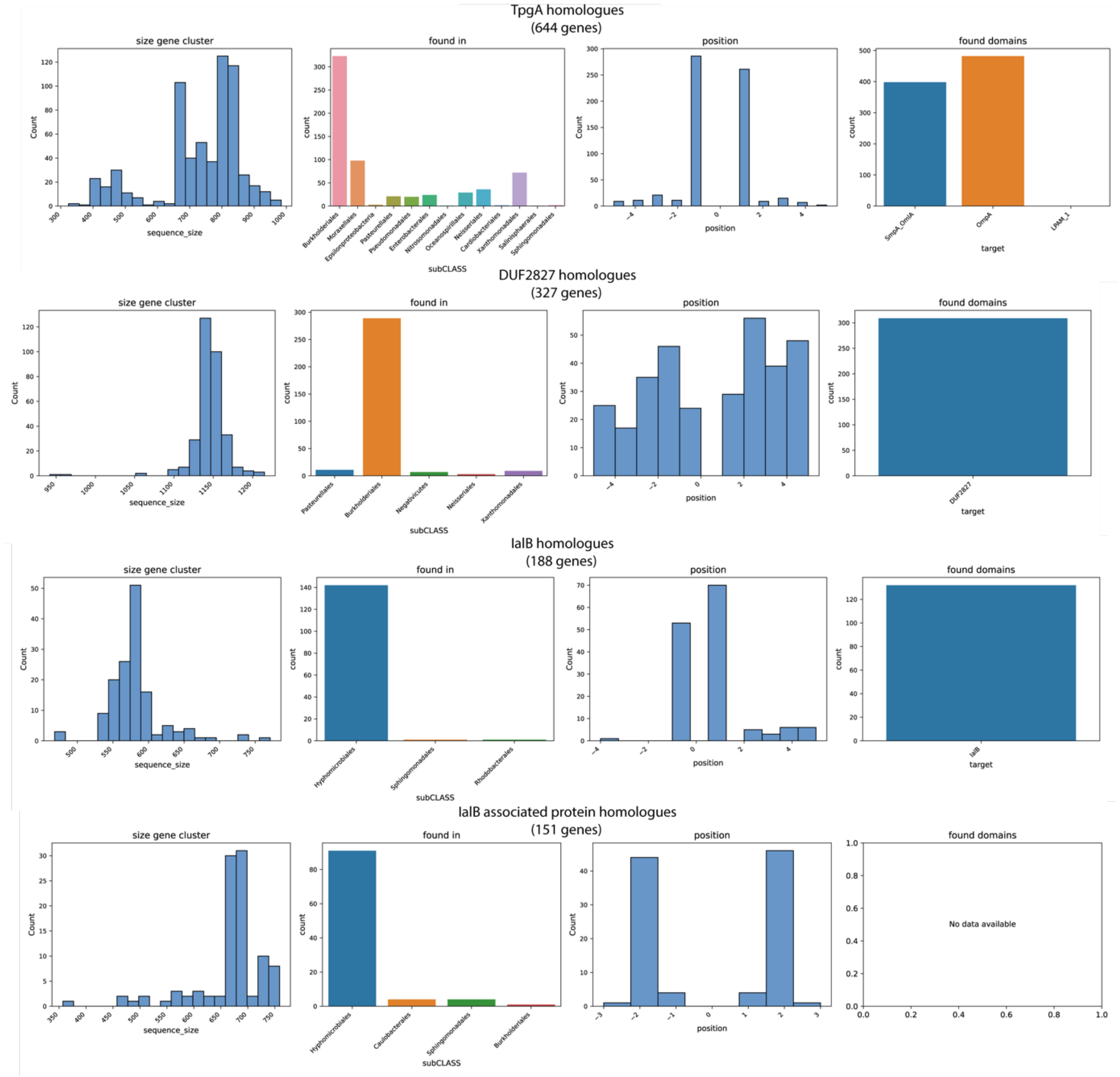

**supplementary Figure S7: Additionnal information concerning the relevant neighbor gene clusters.** For each of the SiliX clusters of TAAs's neighbors is represented an histogram depicting the length distributions (in nucleotides) of the genes found in it, the number of hits per order, an histogram representing the distribution of the relative position to the adhesin (centered on 0) and the associated Pfam domain to each cluster.

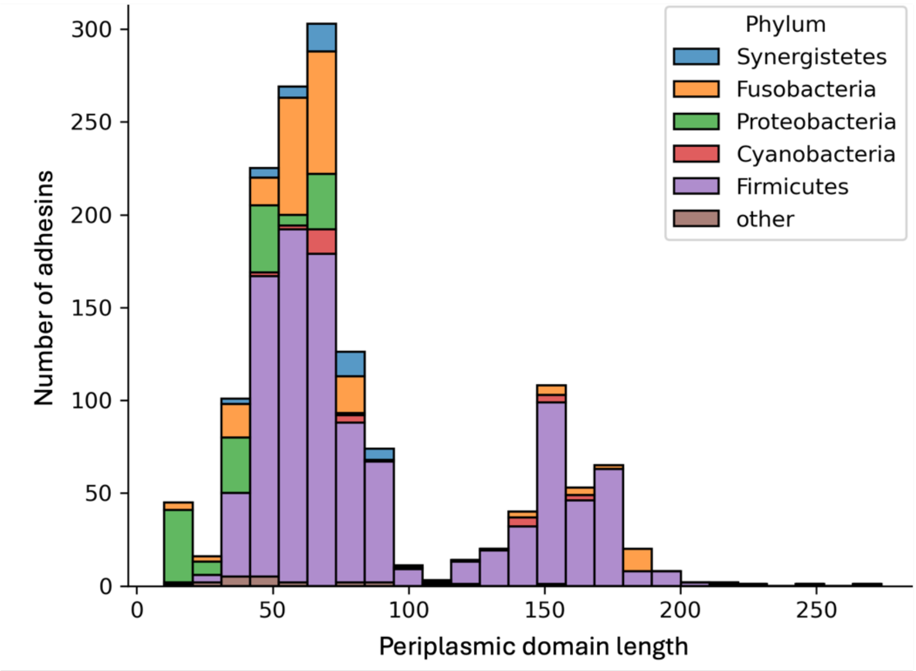

**supplementary Figure S8: Repartition of the identified periplasmic domains based on their length in amino acids.** Histogram representing the distribution of all periplasmic domains identified based on their size and colored by Phylum.

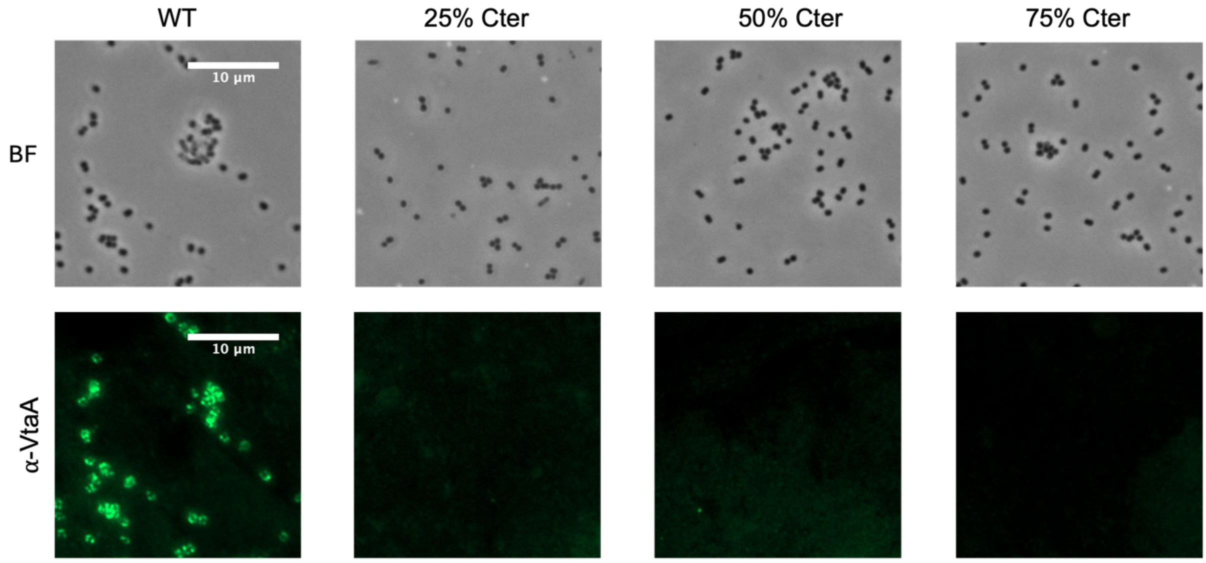

**Supplementary Figure S9: Partial deletion of the VtaA periplasmic domain results in undetectable levels of protein at the cell surface.** Immunofluorescence using an anti-VtaA-head antibody on *V. parvula* WT or with a partial deletion of VtaA periplasmic domain. All images were adjusted to the same contrast values using Fiji (Schindelin et al. 2012). Scale is the same for all images.

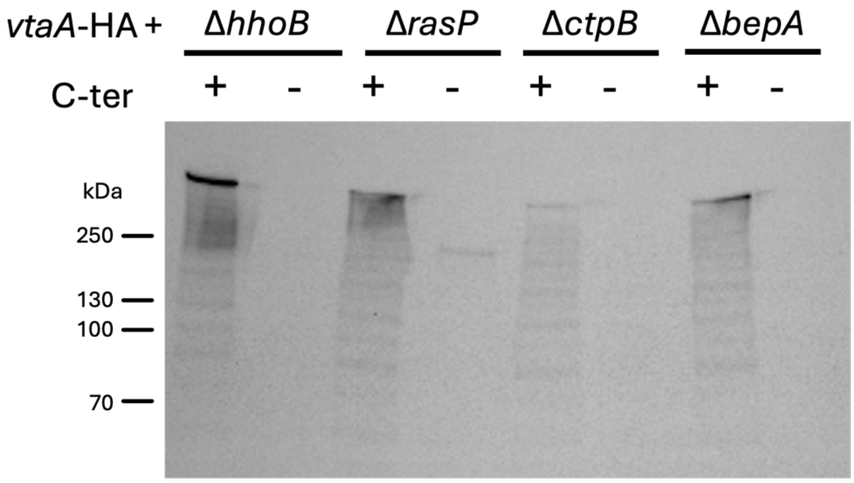

**Supplementary Figure S10: Deletion of putative periplasmic proteases does not impact VtaA stability.** Western-blot with an anti-HA antibody targeting the VtaA-HA-tagged adhesin with or without the periplasmic domain, indicated by the + and - symbols in a background where the gene encoding the indicated protease has been deleted.

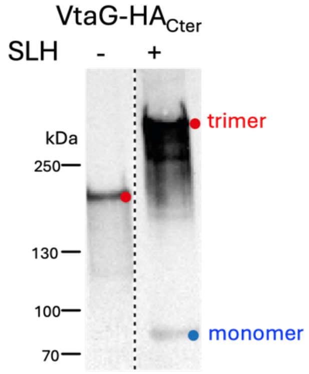

**Supplementary Figure S11: VtaG-HA$_{Cter}$ still forms trimers in absence of the SLH domain.** Western-blot using an anti-HA antibody targeting VtaG with a HA-tag in C-terminal with or without the SLH domain in a Δ*FNLLGLLA_00450-51* background.

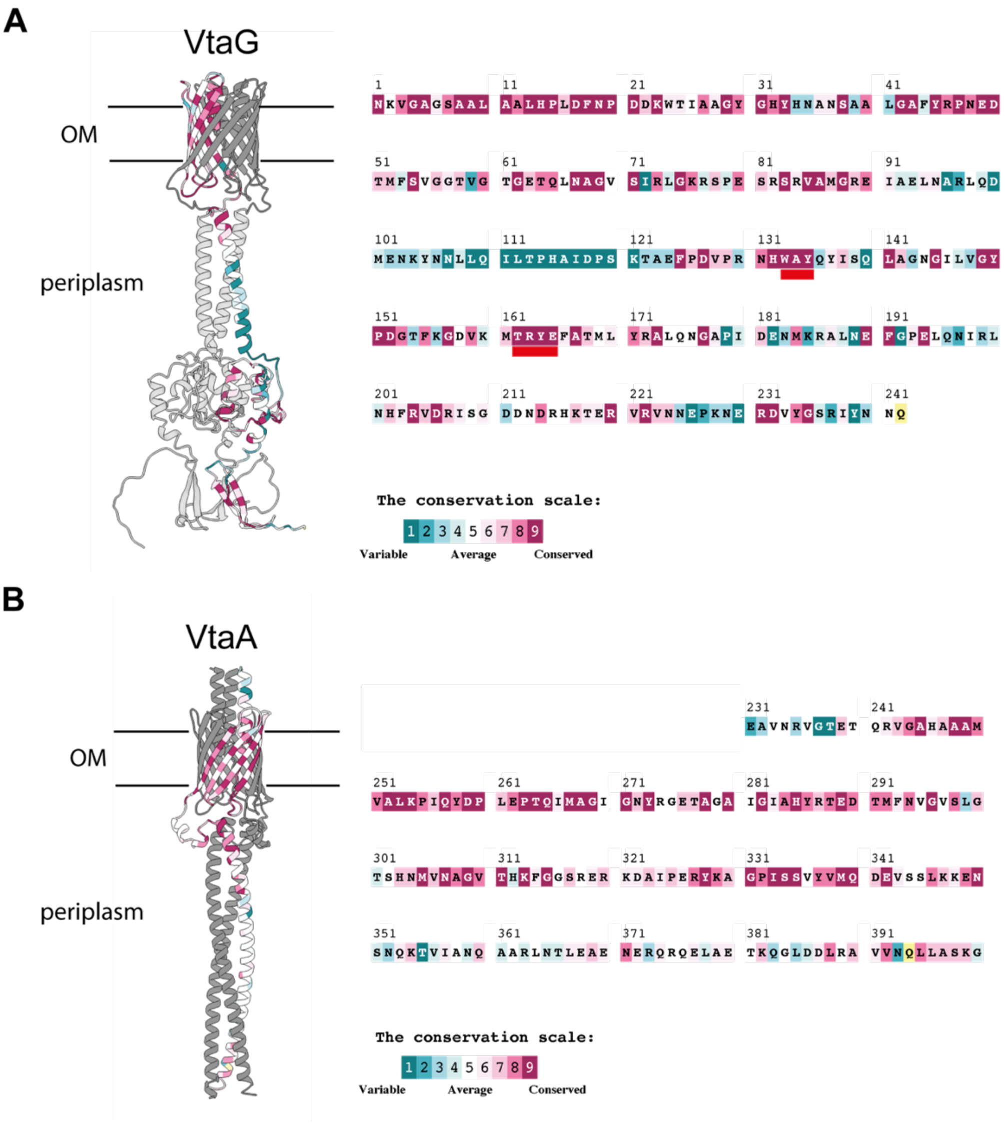

**Supplementary Figure S12: Conservation of the periplasmic domain.** Sequence conservation of VtaG SLH domain (A) and VtaA coiled-coil domain (B) were analyzed using ConSurf (Yariv et al. 2023) with default parameters. Residues are colored by conservation and the conserved WAY and TRYE motifs are underscored in red.

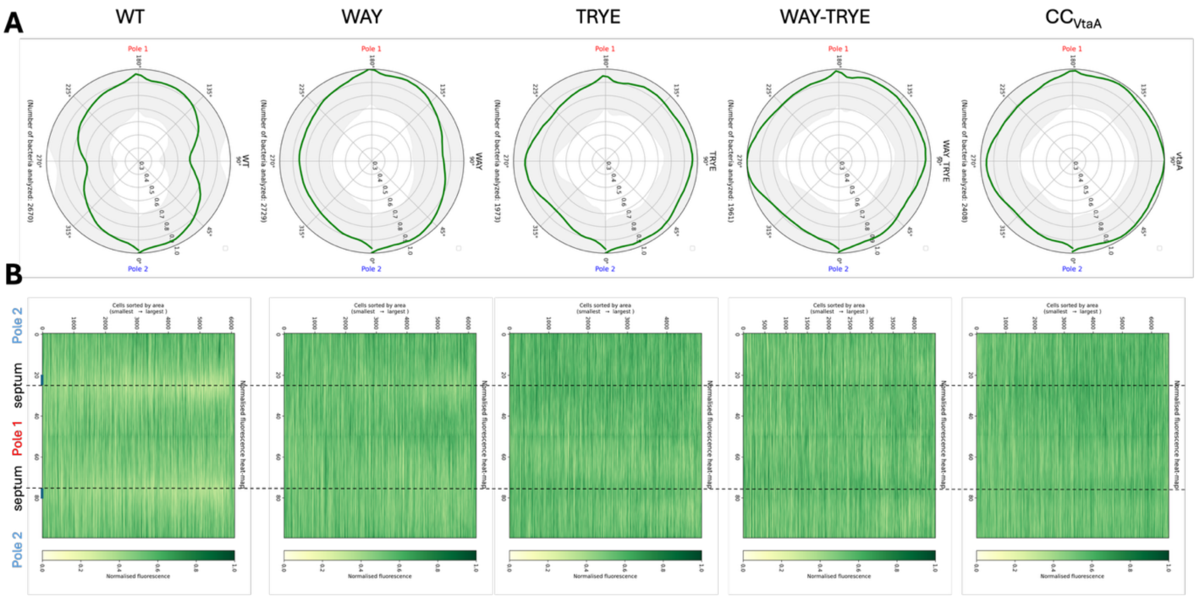

**supplementary Figure S13: VtaG preferentially localizes at the poles in an SLH domain dependant manner.** Immunofluorescence images using an anti-HA antibody on an $HA_{Nter}$-tagged *vtaG* with various modifications of its SLH domain in a Δ*bfvR-S::catP*Δ*vtaF::kanR* background were analysed to measure septum exclusion of VtaG during division A) Polar representation of the average relative normalized fluorescence (green line) and its standard deviation (in gray) for all variants of VtaG. B) Demograph representing the normalized fluorescence for all bacteria analyzed, sorted by size. Each vertical line reprents the fluorescence across the "flattened" perimeter for one bacteria.

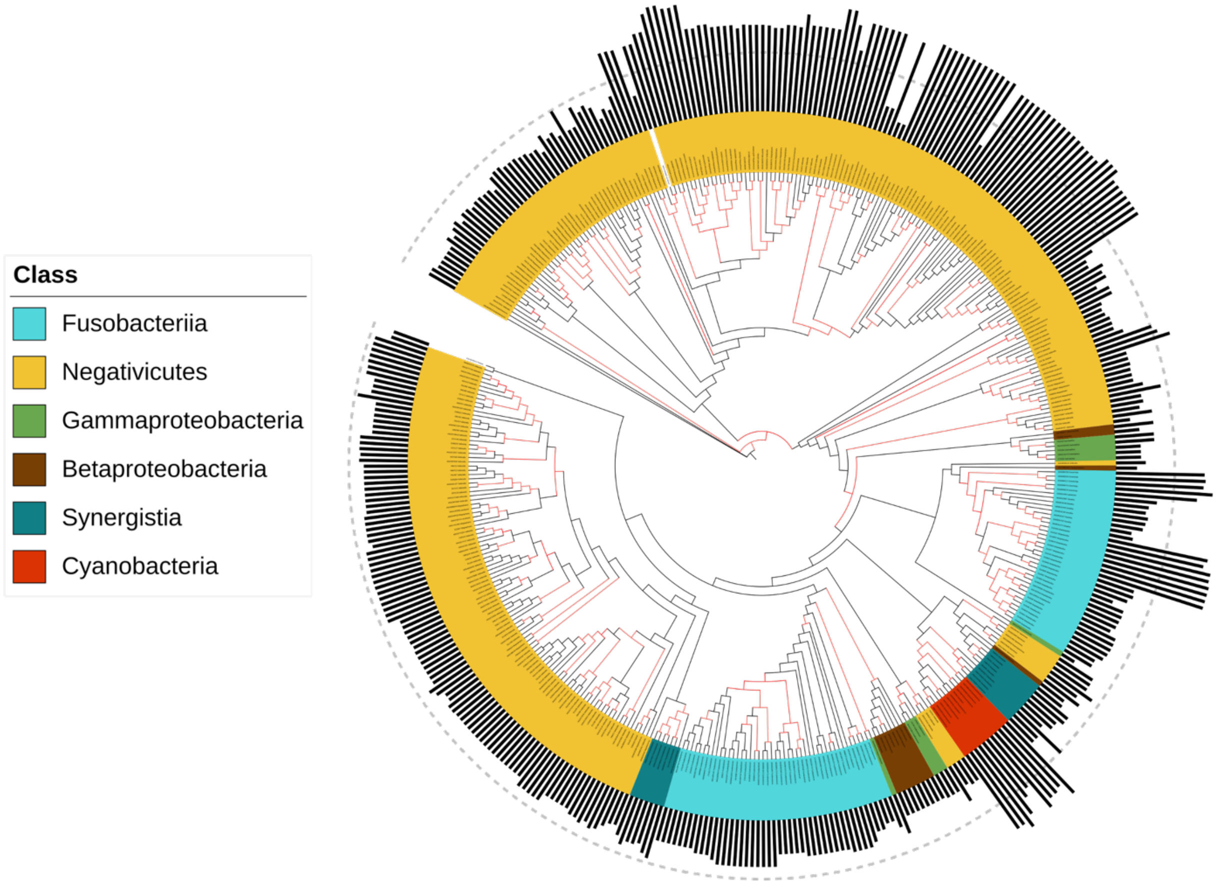

**Supplementary Figure S14: Phylogenetic tree of TAAs containing a periplasmic domain.** The tree was constructed using IQ-TREE based on the TAAs C-terminus sequence including the β-barrel and periplasmic domain sequence. Adhesins are colored by the class of the bacteria they belong. Branches supported by bootstrap (bootstrap above 95) are colored in red, size of the periplasmic domain is indicated by the black bars, with the dotted gray line indicating 100 amino acids size.
